# Hippocampal-prefrontal theta coupling develops as mice become proficient in associative odorant discrimination learning

**DOI:** 10.1101/2021.09.28.462143

**Authors:** Daniel Ramirez-Gordillo, Andrew A. Parra, K. Ulrich Bayer, Diego Restrepo

## Abstract

Learning and memory requires coordinated activity between different regions of the brain. Here we studied the interaction between infralimbic medial prefrontal cortex (mPFC) and hippocampal dorsal CA1 during associative odorant discrimination learning in the male mouse. We found that as the animal learns to discriminate odorants in a go-no go task the coupling of high frequency neural oscillations to the phase of theta oscillations (theta-referenced phase-amplitude coupling or tPAC) changes in a manner that results in divergence between rewarded and unrewarded odorant-elicited changes in the theta-phase referenced power (tPRP) for beta and gamma oscillations. In addition, in the proficient animal there was a decrease in the coordinated oscillatory activity between CA1 and mPFC in the presence of the unrewarded odorant. Furthermore, the changes in tPAC resulted in a marked increase in the accuracy for decoding contextual odorant identity from tPRP when the animal became proficient. Finally, we studied the role of Ca^2+^/calmodulin-dependent protein kinase II α (CaMKIIα), a protein involved in learning and memory, in oscillatory neural processing in this task. We find that the accuracy for decoding the contextual odorant identity from tPRP decreases in CaMKIIα knockout mice and that this accuracy correlates with behavioral performance. These results implicate a role for tPAC and CaMKIIα genotype in olfactory go-no go associative learning in the hippocampal-prefrontal circuit.

**Significance statement:** Coupling of neural oscillations between hippocampal CA1 and medial prefrontal cortex (mPFC) is involved in spatial learning and memory, but the role of oscillation coupling for other learning tasks is not well understood. Here we performed local field potential recording in CA1 and mPFC in mice learning to differentiate rewarded from unrewarded odorants in an associative learning task. We find that odorant-elicited changes in the power of bursts of gamma oscillations at distinct phases of theta oscillations become divergent as the animal becomes proficient allowing decoding of contextual odorant identity. Finally, we find that the accuracy to decode contextual odorant identity decreases in mice deficient for the expression of Ca^2+^/calmodulin-dependent protein kinase II α, a protein involved in synaptic plasticity.

## Introduction

Our lives are enhanced, and our personalities are shaped due to the ability to learn and form memories (Klein et al., 2002). Therefore, it is not surprising that often diseases that affect these abilities are devastating and frequently the individual affected becomes dependent on others. Learning and memory requires coordinated activity between different brain regions (Colgin, 2011; Gordon, 2011; Headley and Paré, 2017; Lisman et al., 2017). Local field potential (LFP) oscillations reflect activity of temporally coordinated neuronal groups providing a reference for spike timing-based codes and gating information transfer between distant brain regions. In theta-referenced phase amplitude coupling (tPAC), the amplitude of the bursts of a faster oscillation is larger within a phase window of a slower carrier wave (Tort et al., 2010). In a previous publication, we characterized tPAC in the olfactory bulb (OB) of mice learning to discriminate odorants in a go-no go associative learning task and we introduced a measure of the magnitude of theta phase-locked high gamma (65-95 Hz) and beta (15- 30 Hz) bursts as a function of time: the theta phase-referenced power (tPRP). We found that tPRP increased for rewarded and decreased for unrewarded odorants in the proficient mouse (Losacco et al., 2020). Furthermore, we showed that contextual odorant identity (is the odorant rewarded?) can be decoded from peak high gamma and peak and trough beta tPRP in animals proficient in odorant discrimination. These findings in the OB, the first relay station in the olfactory system, raised the question whether downstream areas of the brain experience a similar phenomenon.

Here we assessed decoding of contextual odorant identity from oscillatory neural activity in animals learning to discriminate odorants in the go-no go task in two downstream brain areas of the brain that receive input from the OB: CA1 of the hippocampus and medial prefrontal cortex (mPFC) involved in learning of odorant valence and attention to odorants (Cansler et al., 2021; Gourevitch et al., 2010; Li et al., 2017; Martin et al., 2007; Wang et al., 2020). We chose to survey tPAC and tPRP in CA1 and mPFC because optogenetic modulation of interneurons indicates that theta phase-referenced neural activity is involved in memory encoding and retrieval in CA1 (Siegle and Wilson, 2014). We focused on beta and high gamma tPRP because strong directional beta coupling from the OB to the dorsal hippocampus has been shown to be involved in odor processing in the go-no go task (Gourevitch et al., 2010) and because OB spike-high gamma field coherence carries odorant information (Li et al., 2015), and high gamma conveys input from the entorhinal cortex to CA1 (Colgin and Moser, 2010).

Finally, we chose to study odorant decoding in Ca^2+^/calmodulin-dependent protein kinase II α knockout mice (CaMKIIα KO) because of the essential role of this protein in learning and memory (Bayer and Schulman, 2019; Bear et al., 2018; Lisman et al., 2012). CaMKIIα plays a role in long term potentiation (LTP), long-term depression (LTD) and dentate gyrus neurogenesis (Cook et al., 2021; Coultrap et al., 2014; Malinow et al., 1989; Silva et al., 1992b; Suarez-Pereira et al., 2015). Deficiencies in the function of CaMKIIα have been implicated in a range of diseases including schizophrenia, addiction, depression, epilepsy, and neurodevelopmental disorders (Chia et al., 2018; Robison, 2014). CaMKIIα KO are viable and display impaired spatial memory and reduced hippocampal LTP (Silva et al., 1992a; Silva et al., 1992b). Mice heterozygous for CaMKIIα (CaMKIIα Hets) have problems with working memory, increased anxiety, and aggressiveness, characteristic of schizophrenia (Chen et al., 1994; Hasegawa et al., 2009; Matsuo et al., 2009; Yamasaki et al., 2008). Large genome screens have found heterozygous mutations in CaMKIIα in schizophrenia patients (Fromer et al., 2014; Purcell et al., 2014). Furthermore, CaMKIIα Het mice have an immature dentate gyrus (Yamasaki et al., 2008).

## Materials and methods

### Key Resources Table

| REAGENT TYPE | REAGENT or RESOURCE | SOURCE | IDENTIFIER | ADDITIONAL INFORMATION |
| --- | --- | --- | --- | --- |
| Chemical compound, drug | Isoamyl acetate | Sigma-Aldrich | Cat#123-92-2 | Odorant |
| Chemical compound, drug | Ethyl acetate | Sigma-Aldrich | Cat# 141-78-6 | Odorant |
| Chemical compound, drug | Propyl acetate | Sigma-Aldrich | Cat# 109-60-4 | Odorant |
| Chemical compound, drug | Mineral oil | Sigma-Aldrich | Cat# 8042-47-5 | Odorant |
| Chemical compound, drug | Acetophenone | Sigma-Aldrich | Cat# 98-86-2 | Odorant |
| Chemical compound, drug | Ethyl benzoate | Sigma-Aldrich | Cat# 93-89-0 | Odorant |
| Chemical compound, drug | KN93 | MedChemExpress | Cat# HY-15465 | CaMKIIα inhibitor |
| Chemical compound, drug | KN92 | MedChemExpress | Cat# HY-15517 | Inactive derivative of KN93 |
| Other | Nickel-chrome wire | Sandvik | Cat#PX000002 | Tetrode fabrication |
| Other | 16 Channel Electrode Interface Board | Neuralynx | EIB-16 | Tetrode fabrication |
| Software, algorithm | Analysis code | This paper | Available on github and by request | https://github.com/restrepd/drgMaster |
| Strain, strain background | Mouse: CaMKIIα WT | Bayer Lab | Coultrap et al., 2014 | Male mice |
| Strain, strain background | Mouse: CaMKIIα HET | Bayer Lab | Coultrap et al., 2014 | Male mice |
| Strain, strain background | Mouse: CaMKIIα KO | Bayer Lab | Coultrap et al., 2014 | Male mice |
| Software, algorithm | MATLAB_R2018a | Mathworks | RRID: SCR_001622 |  |
| Software, algorithm | Illustrator | Adobe | RRID: SCR_010279 |  |
| Software, algorithm | Photoshop | Adobe | RRID: SCR_014199 |  |

### Animals

Littermate male mice with genotypes of CaMKIIα KO, Het and WT were obtained from heterozygous breeding (Coultrap et al., 2014). Mice were housed in a vivarium with a reversed light cycle of 14/10 h light/dark periods with lights on at 10:00 p.m. Food (Teklad Global Rodent Diet no. 2918; Harlan) was available ad libitum. Access to water was restricted during the behavioral sessions. However, if mice did not obtain ∼1ml of water during the behavioral session, additional water was provided in a dish in the cage (Slotnick and Restrepo, 2005). All mice were weighed daily and received sufficient water to maintain >80% of the weight before water restriction. All experiments were performed according to protocols approved by the University of Colorado Anschutz Medical Campus Institutional Animal Care and Use Committee.

### Surgery and double tetrode implantation

Male mice 2-4 months of age were anesthetized by brief exposure to isoflurane (2.5%) and subsequently anesthesia was maintained with an intraperitoneal injection of ketamine (100 mg/kg) and xylazine (10 mg/kg). The tetrode drive included one optical fiber ferrule for support of an EIB 8 board with 2 tetrodes with four polyamide-coated nichrome wires (diameter 12.5 μm; Sandvik, gold plated to an impedance of 0.2– 0.4MΩ). Tetrodes were connected, and the optic fiber ferrule was glued through an EIB- 8 interface board (Neuralynx). Mice were implanted with two tetrode drives aimed at infralimbic mPFC (+1.94 mm anterior, +0.25 mm lateral, and -3.12 mm below bregma) (Eleore et al., 2011) and the second tetrode drive at the right CA1 layer of the dorsal hippocampus (-3.16 mm posterior, +3.2 mm lateral, and -2 mm below bregma) (Eleore et al., 2011)(Figure 1A). One ground screw was inserted 1 mm posterior from bregma and 1 mm lateral to the midline and sealed to the bone with dental acrylic. Mice were allowed to recover for 1 week before the initiation of the behavioral studies. All behavioral and LFP recording experiments were performed with 2.5-8 month old mice that had undergone double tetrode drive implantation.

**Figure 1.**
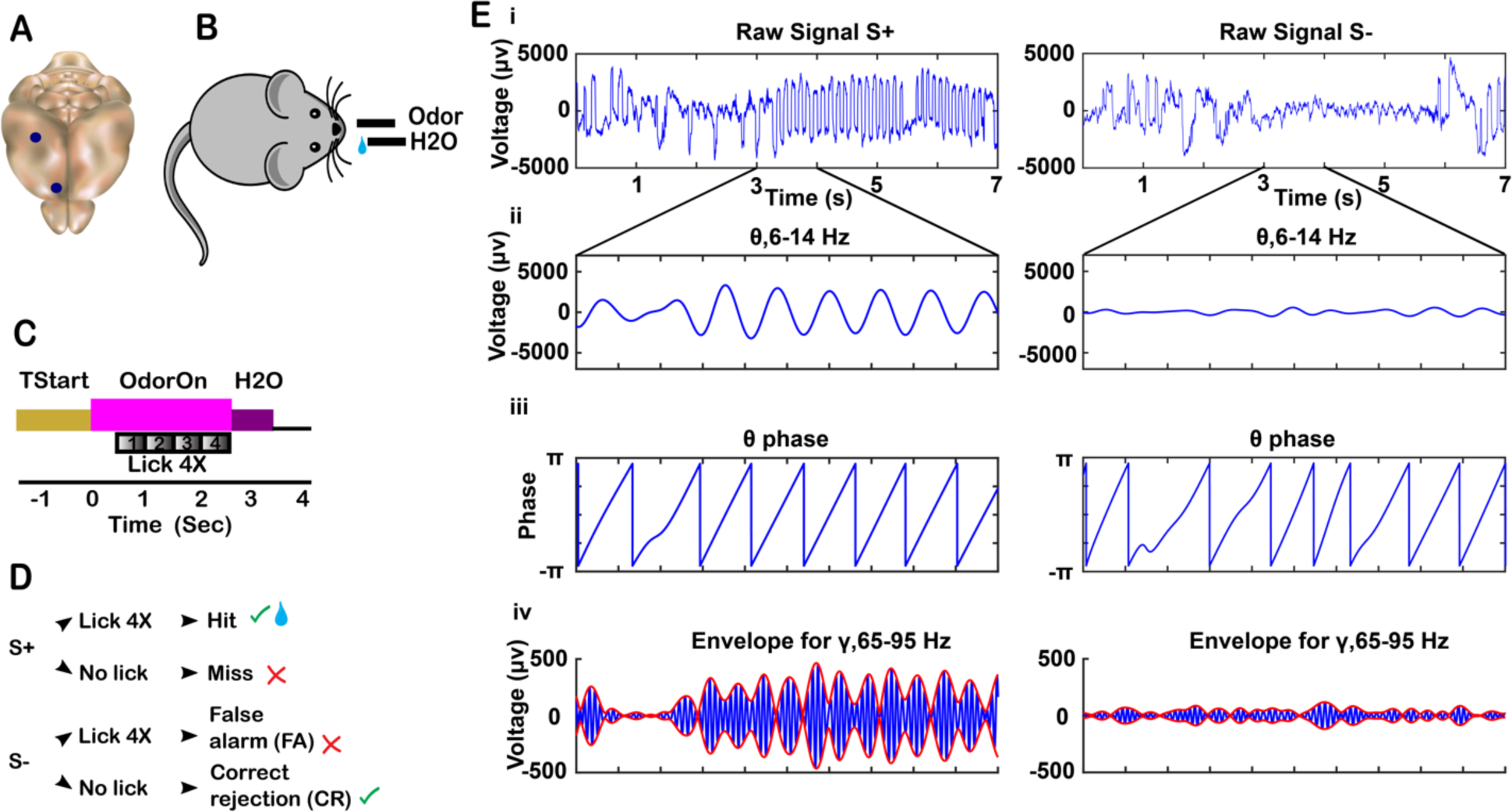
Tetrode location, behavioral task and tPAC analysis. A. Mouse brain showing location of tetrodes in CA1 and mPFC. B. Mouse undergoing the go-no go odor discrimination behavioral task. Mouse self-initiated trials by licking the lixit closing the lick detection circuit. Odorants and water were delivered according to the timeline in C and the decision tree in D. C. Timeline for a single trial. When the animal entered the port and licked at the lixit, it started the trial. TStart (1-1.5 sec) is the time from initiation of the trial to odor delivery. At time 0 the final valve air flow was turned towards the odor port for 2.5 sec resulting in odor onset ∼100 ms after the valve was activated (Losacco et al., 2020). To obtain the water reward for the rewarded odorant (S+) the mouse must lick at least once during each 0.5 s block for four blocks in the response area. If the stimulus was the rewarded odorant and the animal licked, a water reward was delivered (Hit). If the animal did not lick in each 0.5 sec block (Miss), water was not delivered. When the odorant was the unrewarded odorant (S-) the mouse was not rewarded with water regardless of whether it licked in all 0.5 sec blocks (False Alarm, FA) or not (Correct Rejection, CR). D. Decision tree for each trial. Green check mark indicates the correct decision was made. Red “X” mark indicates an incorrect decision was made. Water reward is represented by the water droplet symbol. E. tPAC data analysis for the LFP recorded from CA1. For each electrode, the raw signal collected at 20 kHz (i. rewarded odorant [left], unrewarded odorant right) was filtered into different frequency bands (ii. theta 6-14 Hz). Hilbert transform was used to calculate the theta phase (iii) and the amplitude envelope of higher oscillations such as high gamma (65-95 Hz) (red line in v; blue line is the gamma filtered LFP).

### Go-No-Go Behavioral Task

We used the methods previously described in (Losacco et al., 2020). Briefly, water-restricted mice were required to enter an odor port and lick at the water spout to initiate the release of the odorants 1–1.5 s after the first lick (Figs. 1B,C). Mice were required to lick at least once in four 0.5 s intervals during reinforced odorant delivery (S+) to obtain 10 μl of water. When exposed to the unreinforced odorant (S–), mice refrain from licking for 2 sec. Licking was detected by closing a circuit between the licking spout and the grounded floor of the cage (Slotnick and Restrepo, 2005). The performance of the mice was assessed by calculating correct response to the S+ and S– odorants in 20 trial blocks where 10 S+ and 10 S– odorants were presented at random. Mice were first trained to discriminate between 1% isoamyl acetate (S+) and mineral oil (S-). Once the mice learned to discriminate between isoamyl acetate and mineral oil (percent correct > 80% in two blocks) the odorant pair was switched to the 1% Acetophenone S+ and Ethyl Benzoate S- (APEB) odorant pair. Once the mice learned to discriminate between acetophenone and ethyl benzoate acetate the odor pair was reversed. Ethyl Benzoate became the S+ and Acetophenone the S- (EBAP). When the mice became proficient at the reversal, the odor pair was switched. The same pattern was followed for the rest of the odor pairs (see Table1 for the list of all odorants used). All odorants were obtained from Sigma-Aldrich and were diluted in mineral oil at room temperature. Mice were trained until they performed at 80 percent correct or better in the last two blocks of 20 trials. Figure 2B shows an example of the percent correct odorant discrimination performance per trial. We did not find significant differences in the analysis between odorant pairs.

**Figure 2.**
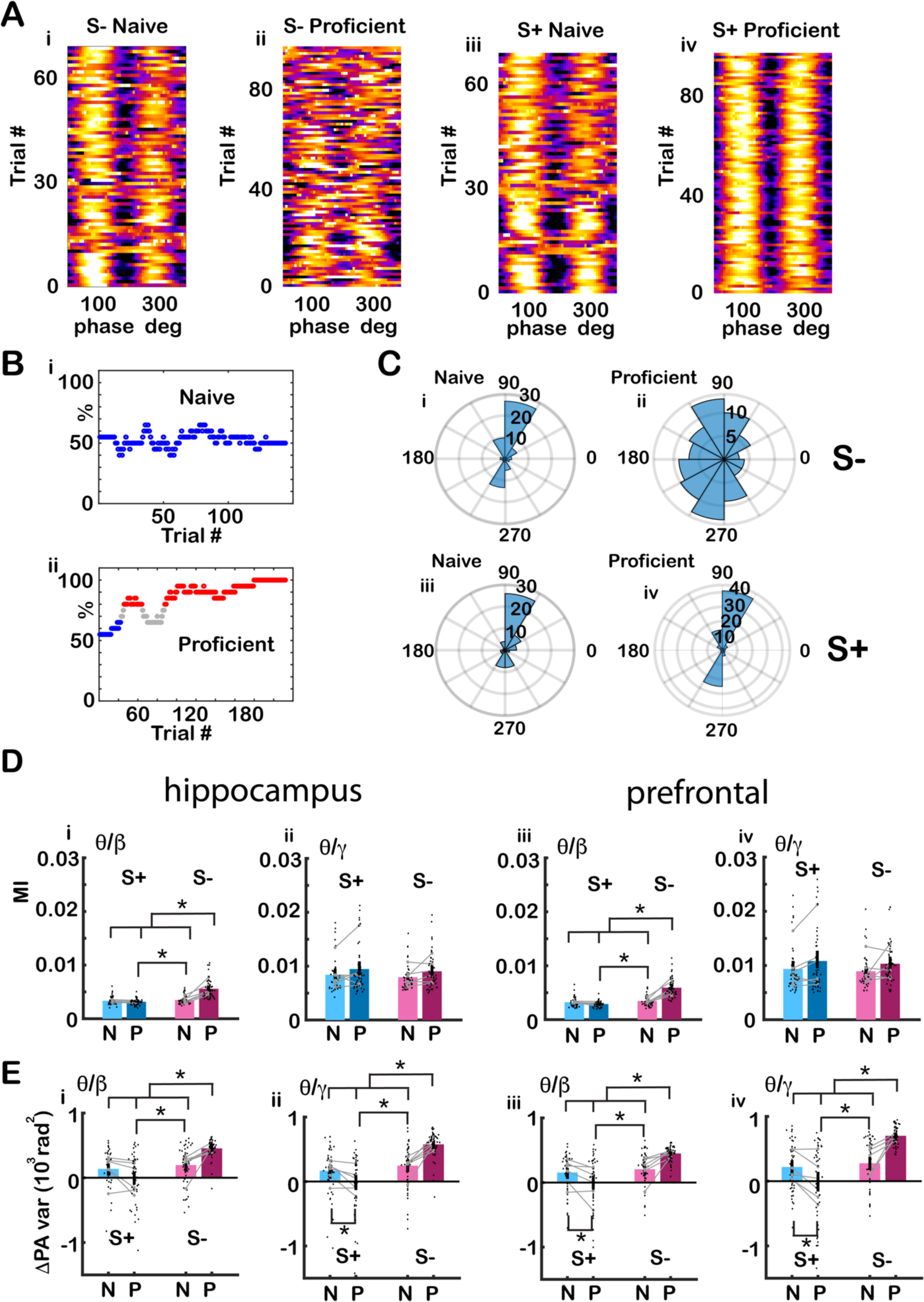
The peak angle variance of tPAC increased for the unrewarded odorant as the animal learned to discriminate odorants. A. Pseudocolor plots showing the phase amplitude relationship for the unrewarded odorant S- (left) and rewarded odorant S+ (right) for two example sessions (naïve and proficient) of the go-no-go task computed for the odorant delivery period (0.5-2.5 sec) for the LFP recorded from CA1. As the animal became proficient differentiating between the rewarded and unrewarded odorants, tPAC peak angle appeared to become more variable for the unrewarded odorant (S-) compared to the rewarded odorant (S+). These pseudocolor plots are all in the same scale. B. Behavioral performance for the two sessions. (i) During the naïve stage, the animal licked for the presentation of the rewarded and unrewarded odorants (performance ∼50%). (ii) During the proficient stage (red points), the animal licked more frequently during the presentation of the rewarded odorant and refrained from licking during the presentation of the unrewarded odorant. Blue = percent correct < 65%, red percent correct >80%. C. Peak angle polar histograms for the examples in A: (i) naïve S- (ii) proficient S- (iii) naïve S+ and (iv) proficient S+. The variance of the peak angle appears to increase for the S- in comparison to the S+ during the proficient stage. D. Bar graphs showing the differences in MI between S+ and S- for the odorant period (0.5-2.5 sec) for hippocampus (i)theta/beta (ii) theta/high gamma and mPFC (iii) theta/beta (iv) theta/high gamma. The black points are per mouse per odorant averages. The grey symbols and lines are per mouse averages. For the beta tPAC for both the hippocampus and mPFC GLM found statistically significant differences for S+ vs. S- and for the interaction between S+ vs. S- and naïve vs. proficient (p<0.001, 188 observations, 184 d.f., F-statistic=51,67.7, p<0.001, 6 mice, 8 odor pairs, Extended Data Fig. 2). E. Bar graphs showing differences for peak angle variance between S+ and S- for the odorant period (0.5-2.5 sec) for hippocampus (i) for theta/beta (ii) theta/high gamma and mPFC (iii) theta/beta (iv) theta/high gamma. The black points are per mouse per odorant averages. The grey symbols and lines are per mouse averages. For both beta and gamma tPAC and for both the hippocampus and mPFC GLM find statistically significant differences for S+ vs. S- and for the interaction between S+ vs. S- and naïve vs. proficient (p<0.001, 188 observations, 184 d.f., F-statistic=19.5-26.4, p<0.001, 6 mice, 8 odor pairs, Extended Data Fig. 2). Asterisks show significant p values (p<pFDR) for post-hoc pairwise tests.

### Local Infusion of KN93 or KN92 into Dorsal CA1

In order to determine the effect of acute local inhibition of CaMKIIα/β in CA1 in mice undergoing the go-no go odorant discrimination task we performed bilateral local infusion of either the CaMKIIα/β inhibitor KN93, or the inactive chemical analogue KN92 into CA1. We used the methods detailed in (Barcomb et al., 2015; Burgdorf et al., 2017). Briefly, a 23-gauge stainless steel guide was attached to an EIB 8 board. A cannular dummy was made from a 27-gauge needle and it was placed in the cannular guide. The device was assembled (cannula + EIB8 board + tetrodes) and was implanted bilaterally into dorsal CA1 (-3.16 mm posterior, +3.2 mm lateral, and -2 mm below bregma) (Eleore et al., 2011). The animals were allowed to recover for a week before behavioral training as described above under the section on the go-no go behavioral task. KN93 and KN92 were obtained from MedChem Express (KN-93 CAS No. 139298-40-1, CAS No. 1431698-47-3) and were initially dissolved in DMSO to a concentration of 10 mM. Shortly before perfusion the drugs were diluted to 10 μM in sterile saline. During drug administration, the guide cannula was fitted with a 27- gauge needle tip that was attached to tubing. The other end of the tubing was attached to a Hamilton syringe. 200 nl of each drug (10 μM) were infused bilaterally using a Razel syringe pump at a rate of 100 nl per minute. The needle was left in place for 4-5 minutes before removing and placing the animal in the olfactometer chamber to perform the go-no-go task. each mouse was randomly assigned to KN92 or KN93. The mice were trained with APEB until they became proficient and then the KN93/KN92 was switched and they were trained with EBAP to proficiency (Table 3).

### Double tetrode recordings

We followed procedures as previously described (Li et al., 2015). The mouse was recorded within the olfactometer chamber with dimensions of 11.6 × 9.7 × 9.4 cm. The EIB-8 boards that recorded signals from the tetrodes were connected to an INTAN RHD2132 16 channel amplifier/A/D converter that interfaced with an RHD2000 USB interface board. Extracellular potentials from the tetrodes were captured, filtered with either a 500 Hz or a 5 kHz low pass filter, and digitized at 20 kHz. Meta data for the behavioral events such as valve opening/closing times and odor identity were recorded through a digital output from the olfactometer. Licks detected by the olfactometer were recorded as an analog signal by the INTAN board.

### tPAC Analysis

As previously described in Losacco et al. (2020), theta-referenced phase amplitude coupling (tPAC) data were processed using the Hilbert transform using a method described by Tort et al. (2010). Briefly, the signal was bandpass filtered with a 20^th^ order Butterworth filter using Matlab’s filtfilt function with zero phase shift to extract LFP in the low frequency oscillation used for phase estimation and the high frequency oscillation used for estimation of the amplitude of the envelope (Figure 1Ei,ii). The Hilbert transform established the theta (6-14 Hz) phase and the envelope for beta (15-30 Hz) and high gamma (65-95 Hz, referred to as gamma) (Figures 1Eiii,iv). To quantify the strength of tPAC, we calculated the modulation index (MI). If tPAC is non-existent, MI = 0, meaning the mean amplitude is distributed uniformly over theta phases, and if tPAC is a delta function MI = 1. MI for signals measured in brain areas such as the hippocampus typically fall between 0–0.03 (Tort et al., 2010).

### tPRP Analysis

As previously described in Losacco et al. (2020), we developed the tPRP approach using custom MATLAB code. tPAC was calculated following the approach used by Tort et al. (2010), as described in tPAC analysis and summarized in Figure 1. Peak and trough theta phases were defined as the peak and trough theta phases for the high frequency oscillations for the S+ trials. A continuous Morlet wavelet transform was used to estimate the power for the high frequency oscillations (Buonviso et al., 2003). tPRP was estimated as the power of the high frequency oscillations measured at the peak or trough of tPAC. The MATLAB code used for data analysis has been deposited to https://github.com/ restrepd/drgMaster. This analysis provides data on which information conveyed by high frequency LFP power is available to a downstream observer restricting observations to a narrow phase window locked to the peak or trough of the theta LFP of the rewarded odorant. Whether tPRP conveyed information on contextual odorant identity was evaluated by a decoding analysis (below).

### Determination of divergence time from ztPRP and lick time courses

We computed a p value with either a ranksum test (for licks) or a t test (for ztPRP) to determine the time when lick rate or ztPRP time courses diverge between rewarded and unrewarded odorant trials in proficient animals. The divergence time was computed as the time point after odorant onset where the p value dropped below 0.005 for >= 1.2 sec after addition of the odorant. As a control, using this criterion on p value traces before odorant application resulted in finding a divergence due to fluctuations in the p values less than 5% of the cases.

### Coherence and Phase Locking Value (PLV) Analysis

In order to quantify the interaction of LFP oscillations from electrodes impaled into different brain regions we computed the complementary values of coherence and phase locking value (PLV). We measured coherence following the method detailed in the supplemental information of (Borjigin et al., 2013). Briefly, coherence between LFP channels is measured by amplitude squared coherence Cxy(f) where f is frequency (mscohere.m in the MATLAB signal toolbox; MathWorks, Inc.). This is a coherence estimate of the input signals x and y using Welch’s averaged, modified periodogram method. The magnitude squared coherence estimate Cxy(f) is a function of frequency with values between 0 and 1 that indicates how well x corresponds to y at each frequency. Furthermore, we quantified phase-locking between the two LFPs by computing PLV following the procedure detailed by Lachaux et al. (1999) using MATLAB code generated by Praneeth Namburi (Namburi, 2021). Briefly, we compute the convolution of each LFP with a complex Gabor wavelet centered at frequency f and then we compute the PLV as the normalized absolute value of the sum of the exponential of the difference in phase multiplied by the imaginary number i. If the phase difference between the two LFPs varies little across the trials, PLV is close to 1; it is close to zero when the relationship between the phases varies randomly across trials.

### Statistical analysis

The statistical analysis was done as described in Losacco et al. (2020) using MATLAB code. Both tPAC and tPRP parameters were calculated separately per electrode (16 electrodes per mouse, 8 hippocampus, 8 mPFC) for all electrodes per mouse. Statistical significance for changes in measured parameters for multivariate factors such as the mouse genotype, naïve vs. proficient and S+ vs. S-, and the interactions of these factors was estimated using generalized linear model (GLM) analysis, with post-hoc tests for all data pairs corrected for multiple comparisons using false discovery rate (Curran-Everett, 2000). The post hoc comparisons between sets of data were performed either with a t-test, or a ranksum test, depending on the result of an Anderson-Darling test of normality. Degrees of freedom and statistical significance have the same meaning in GLM as in analysis of variance and regression (Agresti, 2015). In addition, as a complementary assessment of significant differences (Halsey et al., 2015) we display 95% confidence intervals (CIs) shown in the figures as vertical black lines or shadow boundaries that was estimated by bootstrap analysis of the mean by sampling with replacement 1000 times. In addition, in the Extended Data provided for the figures we include the full result of the GLM analysis and post-hoc t tests or ranksum tests (p<pFDR) as well as the results of a nested ANOVAN taking on account that each odorant pair is run for each mouse. Finally, in order to provide visual information on the distribution of the data for the bar graphs the per odorant per mouse points are spread out along the x axis according to their distribution (this is a violin plot-like display of the data).

### Linear discriminant analysis

Decoding of contextual odorant identity from tPRP values was performed using linear discriminant analysis (LDA) using MATLAB code as described in Losacco et al. (2020). LDA was trained by a leave one out procedure where the algorithm was trained with all trials but one, and the accuracy was assessed by predicting the contextual odorant identity for the trial that was left out. This was repeated for all trials and was performed separately for peak and trough tPRP, and for analysis where the identity of the odorants was shuffled. LDA was performed separately for the naïve and proficient data sets on a per-mouse basis where the input was the tPRP recorded from 16 electrodes.

## Results

### Dual CA1-mPFC tetrode recording in mice undergoing the go-no go olfactory discriminating task

The goal of this study was to determine whether changes in coupled oscillations that occur in the OB as mice learn to discriminate odorants in a go-no go task are observed in downstream areas of the brain (mPFC and hippocampus). These downstream areas receive input originating from the OB and oscillations in the bulb are known to be coupled to the hippocampus and throughout the brain (Martin et al., 2007; Nguyen et al., 2016; Tort et al., 2018). We also wanted to determine whether there was a difference between the CaMKIIα KO, CaMKIIα Hets and the WT mice in theta-referenced phase amplitude coupling (tPAC) that measures the amplitude of high bandwidth beta and gamma oscillations in different phases of slow theta oscillations.

Furthermore, we determined whether the power of high bandwidth oscillations carry different information at the peak and trough of the high bandwidth amplitude phase of the theta oscillation for rewarded odorant trials (beta and gamma tPRP). Evaluating tPRP allows determining whether a downstream observer looking through these two theta phase windows receives different information on the stimulus. Lastly, we determined whether coordinated neural activity changed as the animal learned to discriminate odorants and we asked whether there was a difference in the relationship of oscillations between CA1 and mPFC between the different CaMKIIα genotypes.

In the go-no-go odorant discrimination task, thirsty mice learned to lick on a spout in the presence of a rewarded odorant (S+) to obtain a water reward and refrained from licking in the presence of unrewarded odorant (S-) in a go-no go associative learning task (Losacco et al., 2020). The odorants were presented in pseudorandomized order in the go no-go task (Figure 1A–D). The odorant pairs tested were different volatile compounds (Table 1) and we found no difference in any measurements between odorant pairs. Mice started the trial at will by licking on the lick port. The odorant was delivered at a random time 1–1.5 s after nose poke (the time course for the trial is shown in Figure 1C). The mice learned to either lick a waterspout at least once during each 0.5 s segment in the 2 s response area to obtain a water reward for the rewarded odorant or refrain from licking for the unrewarded odorant. Mice refrained from licking during presentation of the unrewarded odorant likely because of the effort it takes for a sustained 2 sec dry lick period. Behavioral performance was termed naïve or proficient when their performance estimated in a 20-trial window was below 65% for naïve and above 80% for proficient. Mice were trained in the task for sessions of up to 200 trails.

**Table 1.**
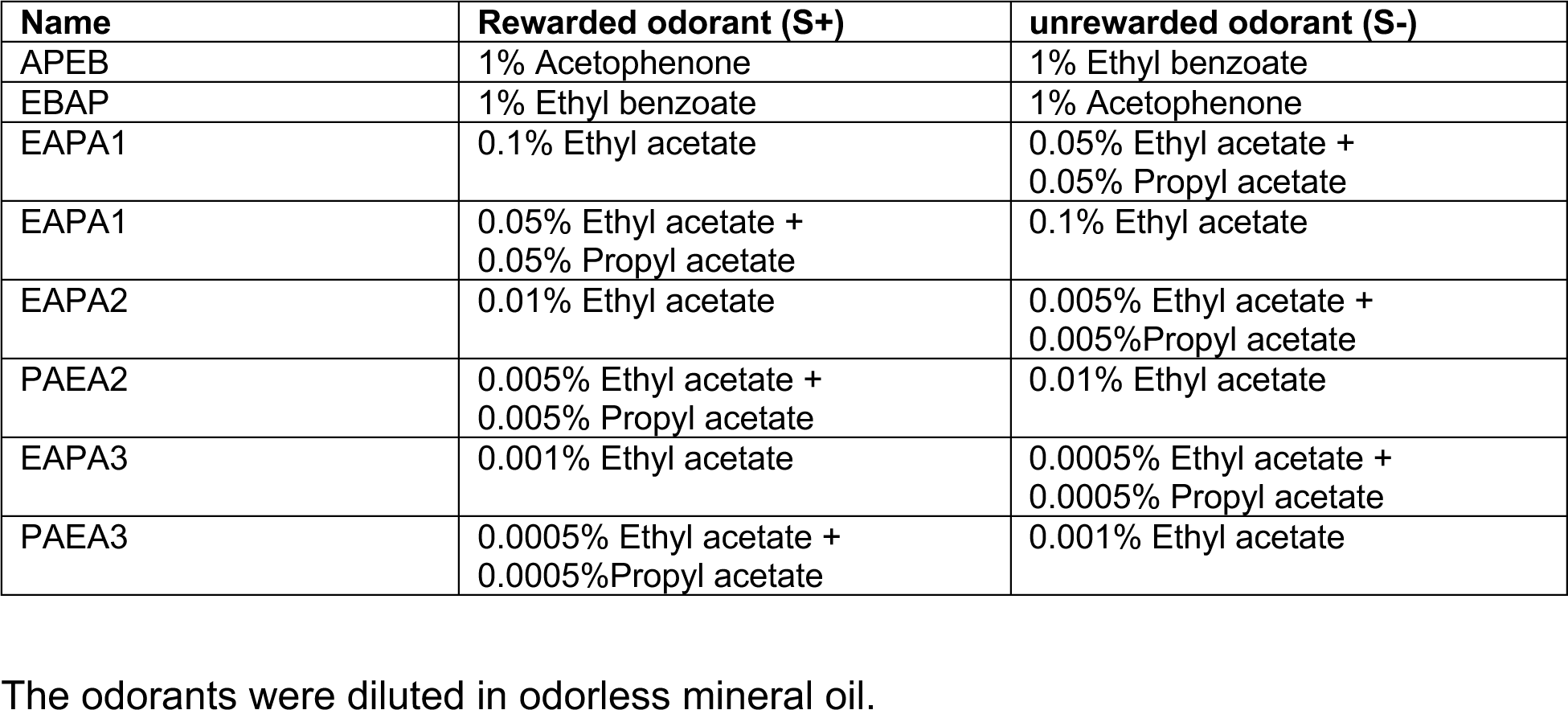
Odorant pairs.

In the last session training ended after the animal achieved two 20-trial blocks of proficient performance; valence was reversed the next day or another odorant was tested. We recorded the LFP using 2 tetrodes (4 electrodes per tetrode) implanted in the dorsal CA1 of the hippocampus and 2 tetrodes in ipsilateral infralimbic medial prefrontal cortex (mPFC), and we analyzed the data to determine whether the different genotypes differ in cross-frequency coupling for naïve or proficient mice (Figure 1E).

The dataset is comprised of 747 recording sessions in 18 mice. The odorant pairs tested were different volatile compounds (or mixtures) whose nomenclature addresses the odorant names (e.g., APEB, see Table 1 for the nomenclature). Table 2 enumerates the total number of sessions per odorant pair, mouse, and experiment.

**Table 2.** Number of sessions and number of mice per odorant pair (Figures 1-13)

| Odorant pair | # of mice | Total sessions |
| --- | --- | --- |
| 1% Acetophenone vs 1% Ethyl benzoate | 6 WT | 22 |
|  | 7 Het | 22 |
|  | 5 KO | 11 |
| 1% Ethyl benzoate vs 1% Acetophenone | 6 WT | 22 |
|  | 7 Het | 55 |
|  | 5 KO | 29 |
| 0.1% Ethyl acetate vs 0.05% ethyl acetate + 0.05% propyl acetate | 6 WT | 15 |
|  | 7 Het | 23 |
|  | 5 KO | 17 |
| 0.05% ethyl acetate + 0.05% propyl acetate vs 0.1% Ethyl acetate | 6 WT | 17 |
|  | 7 Het | 35 |
|  | 5 KO | 37 |
| 0.01% Ethyl acetate vs 0.005% ethyl acetate + 0.005% propyl acetate | 6 WT | 23 |
|  | 7 Het | 52 |
|  | 4 KO | 24 |
| 0.005% ethyl acetate + 0.005% propyl acetate vs 0.01% Ethyl acetate | 6 WT | 31 |
|  | 7 Het | 49 |
|  | 4 KO | 21 |
| 0.001% Ethyl acetate vs 0.0005% ethyl acetate + 0.0005% propyl acetate | 6 WT | 60 |
|  | 7 Het | 64 |
|  | 3 KO | 16 |
| 0.0005% ethyl acetate + 0.0005% propyl acetate vs 0.001% Ethyl acetate | 6 WT | 34 |
|  | 7 Het | 57 |
|  | 3 KO | 13 |

**Table 3.** Number of sessions and number of mice per odorant pair for the drug infusion experiments (**Figure 14**).

| Odorant pair | # of mice | Total sessions |
| --- | --- | --- |
| 1% Acetophenone vs 1% Ethyl benzoate | 5 | 12 |
| 1% Ethyl benzoate vs 1% Acetophenone | 5 | 14 |

We performed tPAC analysis of the LFP recorded in the go no-go behavioral task. tPAC analysis is a cross-frequency coupling analysis to determine whether high frequency oscillation bursts take place at specific phases of low frequency theta oscillations (Tort et al., 2010). tPAC has been reported in brain areas such as the OB, hippocampus and prefrontal cortex (Belluscio et al., 2012; Colgin, 2015; Kaplan et al., 2014; Losacco et al., 2020; Rojas-Libano et al., 2014; Scheffer-Teixeira and Tort, 2016). Figures 1E i–iv show an example of high gamma tPAC for the LFP recorded in CA1. Figure 1E i shows the extracellular LFP sampled for hippocampus at 20 kHz and filtered between 1–750 Hz. The raw signal (Figure 1 E i) was filtered with a 20^th^ order Butterworth filter into different LFP frequency bands (Figures 1E ii, iii, iv) (theta, 6–14 Hz, adapted from (Nguyen Chi et al., 2016) high gamma, 65–95 Hz). We used tPAC analysis (Tort et al., 2010) to evaluate the degree of coupling of the amplitude of the envelope of the beta or high gamma LFP on the phase of the theta LFP. Figure 1E iii shows the theta phase and in Figure 1 E iv shows the envelope for the amplitude of the high gamma LFP, both calculated with the Hilbert transform as detailed in Tort et al. (2010). Figures 1 E ii and iv show that the filtered high gamma LFP changes amplitude in manner that appears coordinated with the theta phase.

### The peak angle variance of tPAC increased for the unrewarded odorant as the animals became proficient differentiating between odorants

We proceeded to ask whether the strength of tPAC, quantified by the modulation index (MI), changes as the animal learns to differentiate odorants in the go-no go task. MI is a measure of how localized high frequency firing is within the phase of theta oscillations (Tort et al 2010). Figures 2A–C illustrate an example of high gamma tPAC recorded in CA1 for S+ and S- odorant trials in two sessions for naïve and proficient mice. The phase amplitude plots for the trials are shown in pseudocolor in Figure 2A, for a WT mouse during the naïve (2Ai-iii) and proficient (2Aii-iv) stages and the percent correct as a function of trial number is shown for the two sessions in Figure 2B. In this example there appears to be an increase in the strength of tPAC when the animal becomes proficient differentiating between the odorants. Furthermore, there was an increase in peak angle variance for the unrewarded odorant as shown by the peak angle polar histograms in Figures 2C i, ii. Figure 2C i shows that for the unrewarded odorant the peak angle was near 90 degrees during the naïve stage while Figure 2C ii shows the peak angle for the unrewarded odorant fluctuated widely during the proficient stage.

This is in contrast with the rewarded odorant for which the peak angle remained near 90 degrees during naïve (Figure 2C iii) and proficient stages (Figure 2C iv).

Figure 2D shows the differences in MI computed per mouse, per odorant pair between S+ and S- for naïve and proficient mice for tPAC for hippocampus (i) beta (ii) high gamma and mPFC (iii) beta (iv) high gamma. For the beta tPAC MI for both the hippocampus and mPFC a generalized linear model (GLM) analysis found statistically significant differences between S+ vs. S- and the interaction between S+ vs. S- and naïve vs. proficient (p<0.001, 188 observations, 184 d.f., F-statistic=51,67.7, p<0.001, 6 mice, 8 odor pairs, Extended Data Fig. 2). GLM does not find significant differences for MI for high gamma tPAC (p>0.05). Figure 2E shows differences for peak angle variance between S+ and S- for naïve and proficient mice for hippocampus tPAC (i) for beta and (ii) high gamma and for mPFC tPAC for (iii) beta and (iv) high gamma. For both beta and gamma tPAC and for both the hippocampus and mPFC GLM found statistically significant differences between S+ vs. S- and the interaction between S+ vs. S- and naïve vs. proficient (p<0.001, 188 observations, 184 d.f., F-statistic=19.5-26.4, p<0.001, 6 mice, 8 odor pairs, Extended Data Fig. 2) and for naïve vs. proficient (p<0.05). Asterisks in all figures show pairwise statistical significance (t test or ranksum test, p< pFDR, p value for significance corrected for multiple comparisons using the false discovery rate (Curran-Everett, 2000)).

Overall, we found an increase in peak angle variance for the unrewarded odorant as the animal became proficient. We also found small, but significant changes in the strength of beta tPAC that differed between the S+ and S- odorants as the animal learned to discriminate odorants.

### The odorant-elicited change in the theta phase-referenced beta and gamma power became negative for the unrewarded odorant as the mice became proficient discriminating between odorants

Wavelet power referenced to theta phase (peak or trough) was determined to evaluate whether it changes as the animal learns. This analysis is referred to as theta phase-referenced power (tPRP) (Losacco et al., 2020). Figure 3A shows examples for CA1 of the time course during the trial for the average wavelet broadband LFP spectrograms for 30 trials during naïve S+ (i), 27 trials during naïve S- (ii) 84 trials during proficient S+ and 84 trials during proficient S- (iv). After odorant onset there was an increase in broadband power for the S+ odorant and a decrease in power for the S- odorant as the animal became proficient.

**Figure 3.**
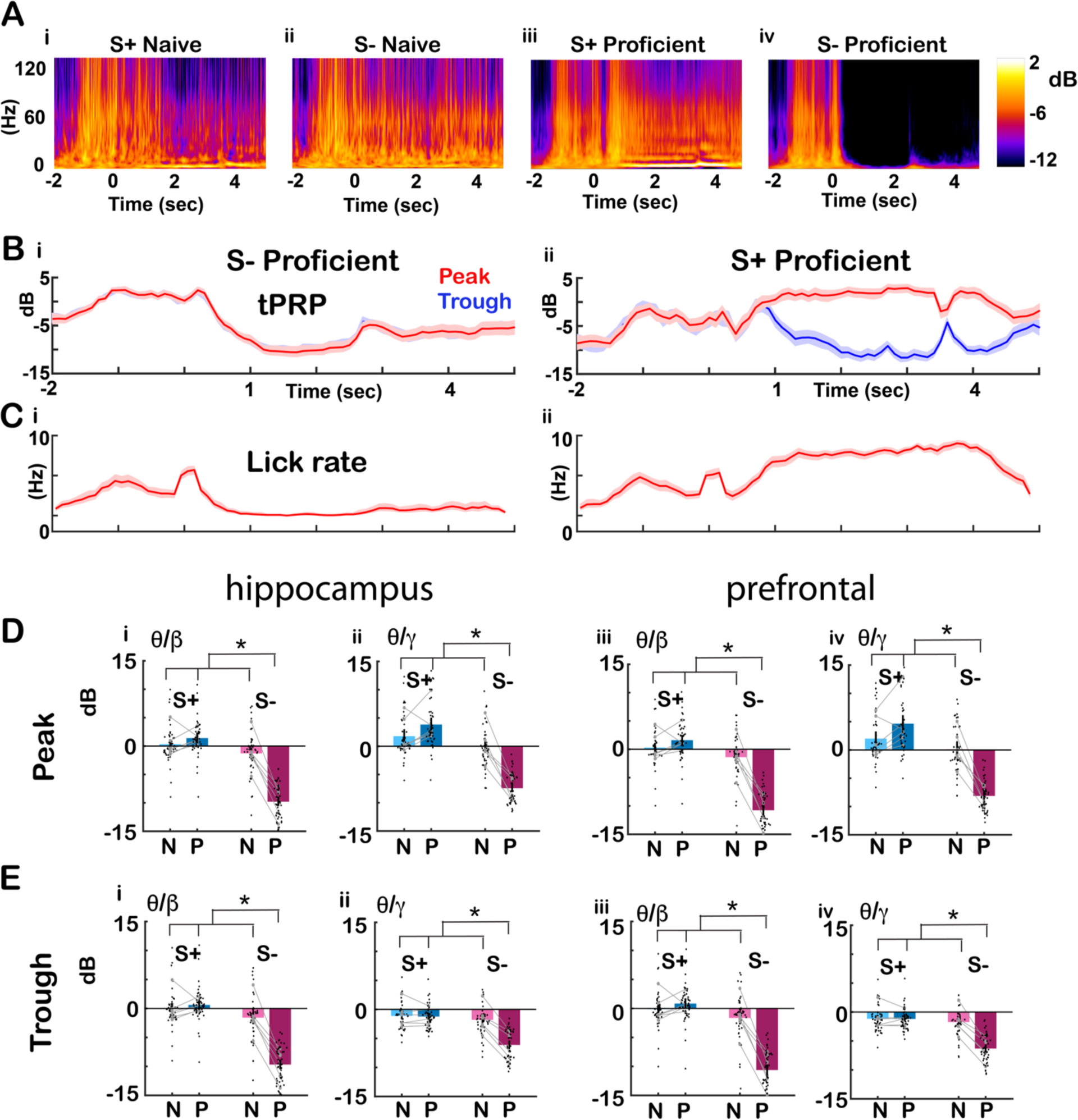
Changes in phase referenced power as the animal became proficient. A. Examples of the time course per trial for the average wavelet broadband LFP spectrogram for (i) S+ naïve (30 trials) (ii) S- naïve (27 trials) (iii) S+ proficient (84 trials) (iv) S- proficient (84 trials). Pseudocolor scale for LFP power is shown in dB. The LFP was recorded from CA1. B. Average gamma wavelet power referenced to theta peak and through for the same trials as shown in A (i) S- proficient (ii) S+ proficient, red: peak, blue: through, shadow: confidence interval. C. Lick rate for the same trials (i) S- proficient (ii) S+ proficient. D and E. Summary bar graphs showing that peak/trough tPRP for the odorant period (0.5-2.5 sec after the odorant is diverted to the odor port) decreases for S- as the mice become proficient. Black points are per mouse per odorant averages. The grey symbols and lines are per mouse averages. D: peak, E: through, (i) theta/beta and (ii) theta/gamma hippocampus (iii) theta/beta and (iv) theta/gamma prefrontal cortex. GLM found statistically significant differences for S+ vs. S- and for the interaction between S+ vs. S- and naïve vs. proficient (p<0.001, 376 observations, 368 d.f., F-statistic=72.7-103, p<0.001, 6 mice, 8 odor pairs, Extended Data Fig. 3). Asterisks show significant p values (p<pFDR) for post-hoc pairwise tests.

Figure 3B shows for this example the time course for the LFP recorded in CA1 during the trial for the average high gamma tPRP referenced to the peak (red) or trough (blue) of theta for the proficient trials in Figure 3A. For the rewarded (S+) odorant the power increased for the peak and decreased for trough after addition of the odorant while for the unrewarded odorant both peak and trough tPRPs decreased. Figure 3C shows that, as expected, there was an increase in the lick rate for the rewarded odorant and a decrease for the unrewarded odorant. Finally, Figures 3D,E show a per mouse per odorant pair analysis that indicated that tPRP became negative for S- as the mice became proficient. E: peak, F: through, (i) beta and (ii) gamma hippocampus (iii) beta and (iv) gamma mPFC. GLM found statistically significant differences for tPRP between S+ vs. S- and the interaction between S+ vs. S- and naïve vs. proficient (p<0.001, 376 observations, 368 d.f., F-statistic=72.7-103, p<0.001, 6 mice, 8 odor pairs, Extended Data Fig. 3).

In addition, we asked whether odorant-induced changes in peak tPRP changed when odorant valence was reversed. Panels A i and ii of Figure 4 show an example for CA1 LFP of the time course for the peak (red) and trough (blue) high gamma tPRP when the mouse was proficient for a forward session where the rewarded odorant (S+) was acetophenone (AP) and the unrewarded odorant (S-) was ethyl benzoate (EB). Panels A iii and iv of Figure 4 show that when the valence of the odorant was reversed (AP was S- and EB was S+) the response to EB resembled the response to AP in the forward sessions (compare panels A ii and iv of Figure 4) indicating that the response is a response to the contextual identity of the odorant (is it rewarded?) as opposed to the chemical identity of the odorant. Figure 4 (B,C) shows a summary bar graph analysis per mouse per odorant pair of all reversal experiments indicating that for the proficient mouse the average peak and trough tPRP decreases for the unrewarded S- odorant regardless of the identity of the odorant. For high gamma tPRP GLM found statistically significant differences for S+ vs. S- and peak vs. trough (p<0.001, 188 observations, 180 d.f., F-statistic=61.8, p<0.001, 6 mice, 8 odor pairs, Extended Data Fig. 4) and does not find a difference between forward and reverse sessions indicating that indeed the high gamma tPRP responds to the contextual odorant identity. For beta tPRP GLM found a statistically significant difference for S+ vs. S- (p<0.001, 188 observations, 180 d.f., F-statistic=87, p<0.001, 6 mice, 8 odor pairs, Extended Data Fig. 4) and interestingly, GLM does find a small statistically significant differences between forward vs. reverse (p<0.05) for beta tPRP. Although this is a relatively small difference, this indicates that beta tPRP is not exclusively responsive to contextual odorant identity and that these brain regions may also encode information on the chemical identity of the odorant.

**Figure 4.**
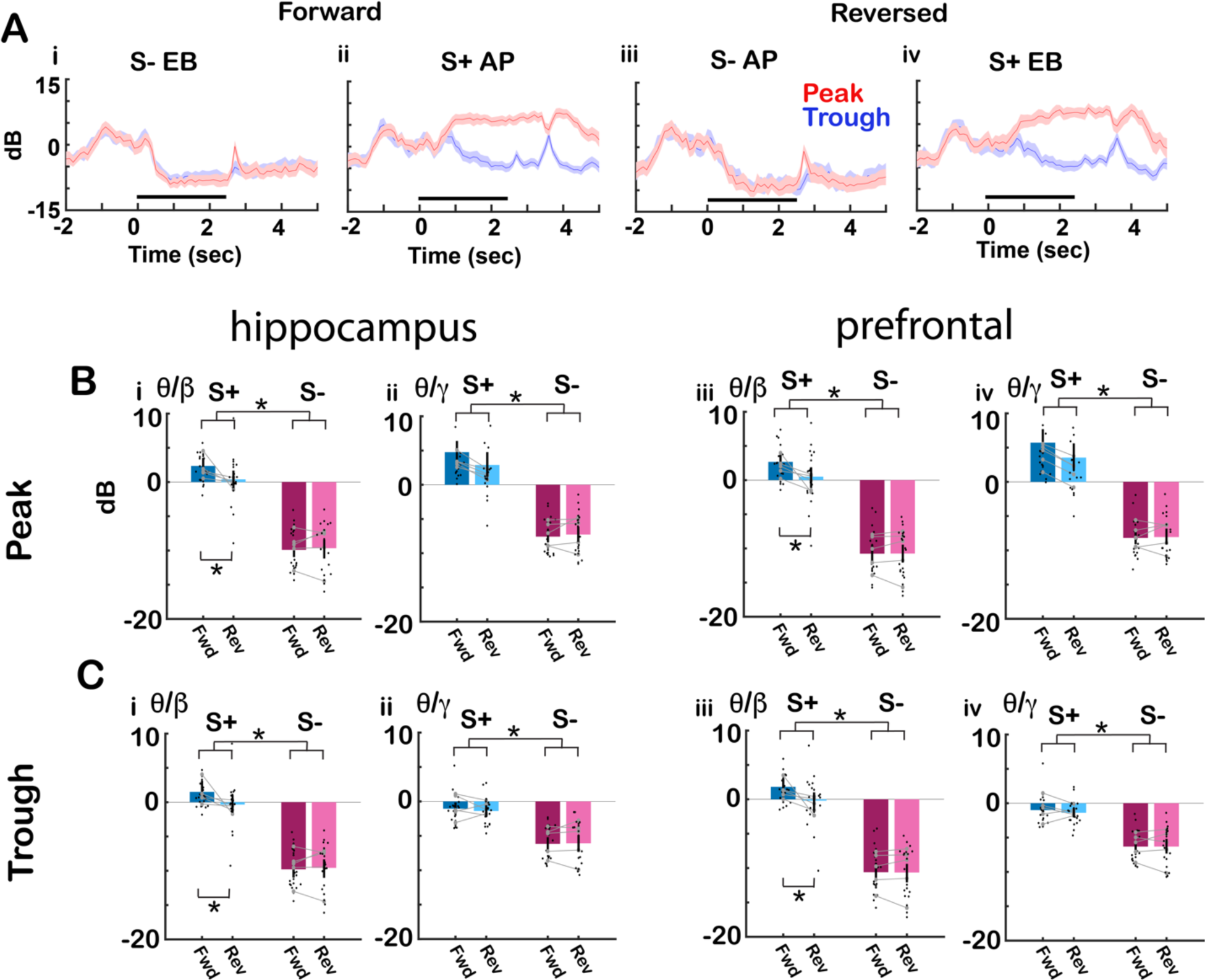
Effect of reversal of odorant valence on the phase referenced power. (A) Average gamma wavelet power referenced to theta peak and through for the same trials when the mouse was proficient for the CA1 LFP example shown in Figure 3A. The gamma tPRP is normalized by subtracting the power before odorant application. (i,ii) Forward: (i) S- (ethyl benzoate, EB) (ii) S+ (acetophenone, AP); (iii,iv) Reversed: (iii) S- (acetophenone, AP) (iv) S+ (ethyl benzoate, EB), red: peak, blue: through, shadow; confidence interval. (B,C) Summary bar graphs showing that the average peak tPRP for the odorant period (0.5-2.5 sec after the odorant is diverted to the odor port) decreases for S- upon odorant application regardless of the identity of the odorant. Black points are per mouse per odorant averages. The grey symbols and lines are per mouse averages. B: peak, C: through, (i) theta/beta and (ii) theta/gamma hippocampus (iii) theta/beta and (iv) theta/gamma prefrontal cortex. For beta tPRP GLM found statistically significant differences for forward vs. reverse (p<0.05) and S+ vs. S- (p<0.001, 188 observations, 180 d.f., F-statistic=87, p<0.001, 6 mice, 8 odor pairs, Extended Data Fig. 4). For gamma tPRP GLM found statistically significant differences for S+ vs. S- and peak vs. trough (p<0.001, 188 observations, 180 d.f., F-statistic=61.8, p<0.001, 6 mice, 8 odor pairs, Extended Data Fig. 4). Asterisks show significant p values (p<pFDR) for post-hoc pairwise tests.

### The accuracy for decoding the contextual identity of the odorants from theta phase referenced power increased when the mice became proficient

We proceeded to determine whether we could decode contextual odorant identity from tPRP. Decoding was performed using a linear discriminant analysis (LDA) to set a decision boundary hyper-plane between binary stimulus classes (S+ versus S-) (Vizcay et al., 2015). LDA was trained with tPRP from each electrode for each mouse (8 electrodes in the hippocampus and 8 electrodes in the mPFC per mouse) for all trials except one (the training dataset) and then the tPRP from the missing trial (test data) was classified as S+ or S-. This training was performed separately at both naïve and proficient learning stages. As a control we shuffled the identity of trials in the training set.

Figure 5A shows an example of the time course during the trial for the decoding accuracy for one mouse for the LDA trained using CA1 tPRP for the EAPA odor pair for (i) naïve stage beta, (ii) proficient stage beta, (iii) naïve stage gamma, (iv) proficient stage gamma. For the naïve animal decoding accuracy increased slowly after the addition of the odorant, and increased further after the animal received the reward (Figures 5Ai,iii). When the animal became proficient decoding accuracy increased rapidly beyond 80% after addition of the odorant (Figures 5Aii,iv). For high gamma tPRP the accuracy was higher for the proficient animal for peak-referenced tPRP compared to trough-referenced tPRP (Figure 5Aiv). Figures 5 B and C show mean bar graphs for the mean accuracy for decoding contextual odorant identity calculated for the last second of the odorant epoch (1.5 to 2.5 sec after diverting the odorant towards the mouse) for shuffled, naïve, and proficient. Figure 5B shows the accuracy for the peak tPRP and Figure 5C shows the accuracy for trough tPRP. For all conditions the accuracy is significantly higher for the proficient stage compared to naïve or shuffled. GLM analysis found for both CA1 and mPFC statistically significant differences between naïve vs. proficient and shuffled vs. proficient for both beta and gamma tPRP (p<0.001, 756 observations, 744 d.f., F-statistic=25.1, p<0.001, 6 mice, 8 odor pairs, Extended Data, Fig. 5) and for gamma tPRP GLM found statistically significant differences between peak and trough (p<0.001, 756 observations, 744 d.f., F-statistic=25.2, p<0.001, 6 mice, 8 odor pairs, Extended Data Fig. 5).

**Figure 5.**
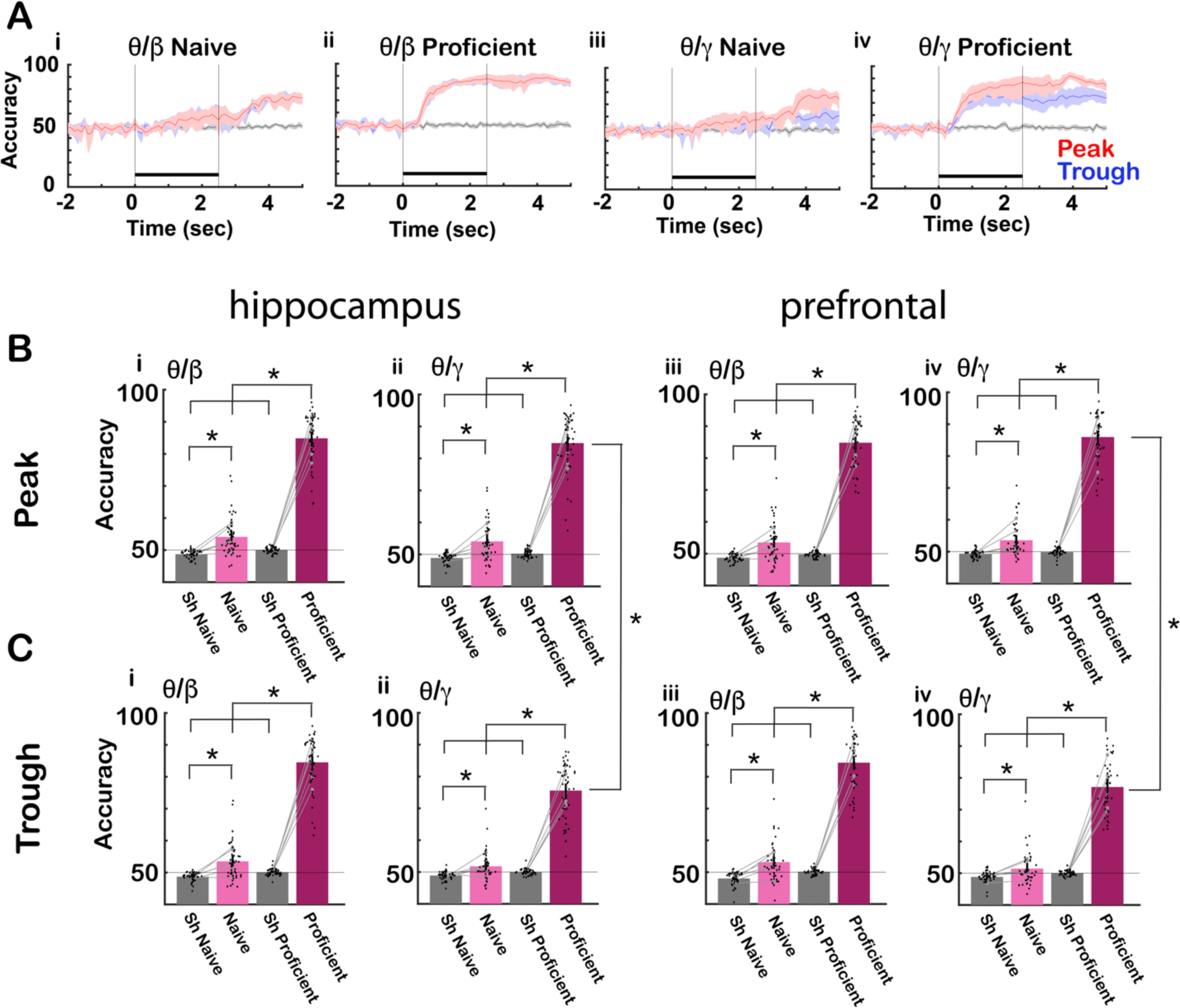
Linear discriminant analysis for decoding the contextual odorant identity from tPRP. A. Example for one mouse for the time course for the accuracy of odorant identity decoding by a linear discriminant analysis algorithm trained using tPRP calculated from CA1 LFP for the EAPA odor pair (i) naïve theta/beta, (ii) proficient theta/beta, (iii) naïve theta/gamma, (iv) proficient theta/gamma red: peak, blue: through, black: shuffled, shadow: confidence interval, black bar: odorant application. B and C. Bar graphs showing the differences in decoding accuracy between shuffled, naïve, and proficient. B: Accuracy for peak tPRP for (i) theta/beta in the hippocampus, (ii) theta/gamma in the hippocampus, (iii) theta/beta in mPFC, (iv) theta/gamma in mPFC. C: Accuracy for through for (i) theta/beta in the hippocampus, (ii) theta/gamma in the hippocampus, (iii) theta/beta in mPFC, (iv) theta/gamma mPFC. The bars show the average accuracy, and the points are the accuracy per mouse per odor pair. The vertical bars show the confidence interval. The grey symbols and lines are per mouse averages. For beta and gamma tPRP for both prefrontal and hippocampus LDA GLM found statistically significant differences for naïve vs. proficient and shuffled vs. proficient (p<0.001, 380 observations, 372 d.f., F-statistic=355-494, p<0.001, 6 mice, 8 odor pairs, Extended Data Fig. 5). For gamma tPRP for both prefrontal and hippocampus LDA GLM found statistically significant differences between peak and trough (p<0.05, 380 observations, 372 d.f., F-statistic=355-494, p<0.001, 6 mice, 8 odor pairs, Extended Data Fig. 5). Asterisks show significant p values (p<pFDR) for post-hoc pairwise tests.

### Theta phase referenced power diverges between rewarded and unrewarded trials before divergence of lick behavior

Theta oscillations in mPFC are phase locked with licks in rats consuming liquid sucrose rewards and this theta range activity has been postulated to encode for the value of consumed fluids (Amarante et al., 2017; Amarante and Laubach, 2021). We proceeded to analyze the relationship between the tPRP time course in CA1 and mPFC and the time course for licks in the go-no go task. Figure 6A shows that licks are phase locked to the theta LFP for an example of licks aligned with theta LFP recorded from CA1 for a hit trial. Figure 6B shows the lick traces for the rewarded and unrewarded odorants for a proficient animal engaged in the go-no go task with the APEB odorant pair. For the rewarded odorant the animal licked continuously for several seconds after odorant application while for the unrewarded odorant the animal stoped licking shortly after the odorant was delivered. The top panel shows that the p value calculated using a ranksum test of the difference in binary lick recordings between S+ and S- trials decreased sharply shortly after addition of the odorant reflecting divergence in lick behavior between the rewarded and unrewarded odorants. Figure 6C shows the mean lick rate for five mice undergoing the go-no go task for the APEB odorant pair.

**Figure 6.**
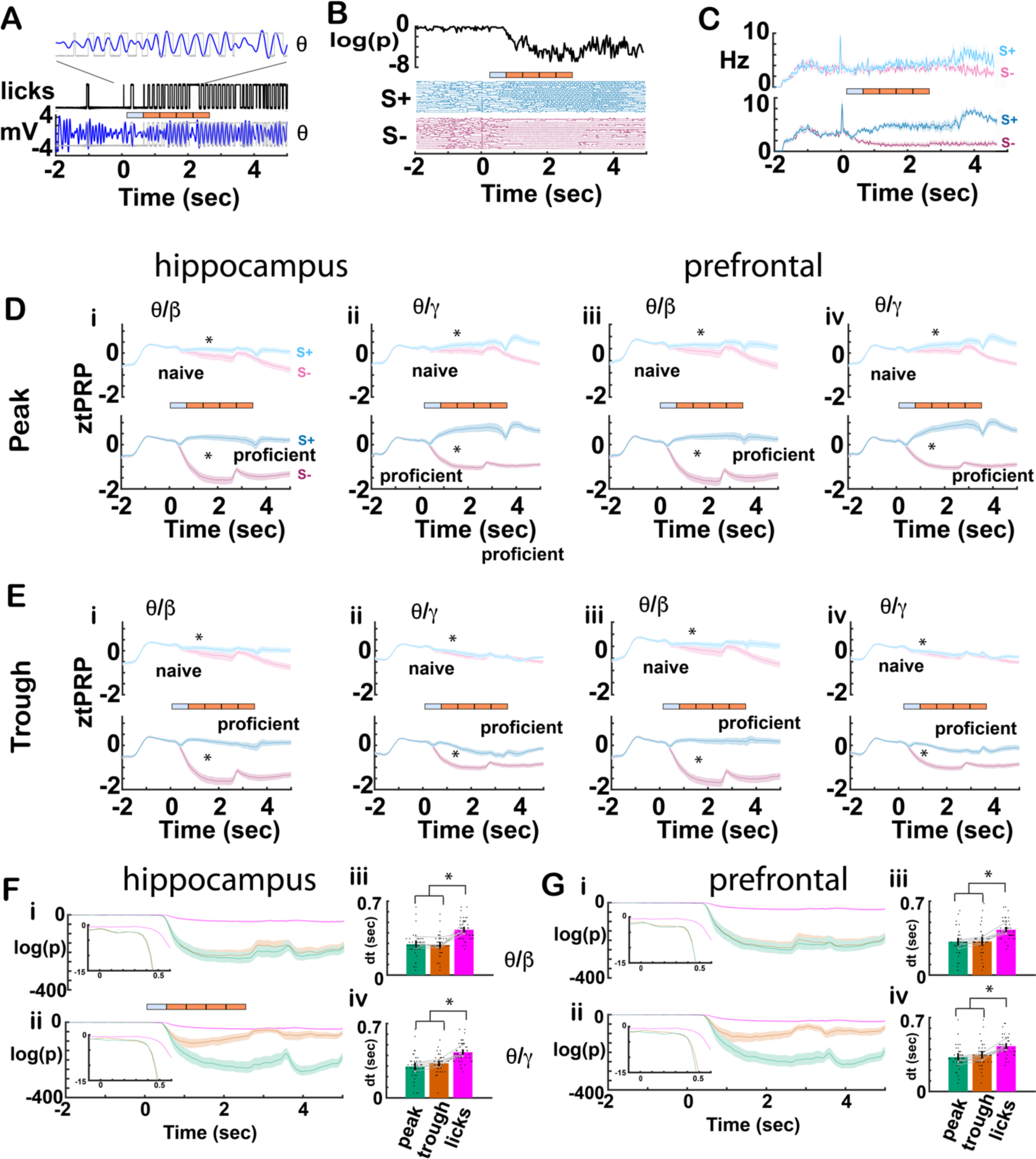
Comparison of divergence between rewarded and unrewarded trial time courses for lick rate and tPRP. A. Example illustrating the relationship between licks and theta CA1 LFP for a Hit trial. The lick signal is binary with licks represented by an increase in voltage. The bar shows the 2.5 sec odorant period divided in 0.5 sec response areas. The mouse must lick at least once in each of the last four response areas to obtain a reward. B. Lick time course for a proficient mouse for 18 S+ and 18 S- trials (bottom) and for the p value for the ranksum test comparing the S+ and S- binary lick traces (calculated in 33 msec time bins). C. Average lick rate time course for the APEB odorant pair (±CI) for naïve and proficient mice for the rewarded (S+) and unrewarded (S-) odorants (average of five mice). D and E. Time course for the average tPRP. The tPRP is z normalized by subtracting the mean tPRPfrom -2 to 0 sec and dividing by the standard deviation in this period. The bounded line shows the average ztPRP bounded by the CI calculated from ztPRP traces per odorant per mouse (6 mice, 8 odor pairs). D is the peak ztPRP and E is the trough ztPRP. i, hippocampus theta/beta, ii, hippocampus theta/gamma, iii, mPFC theta/beta and iv, mPFC theta/gamma. Cyan is S+ and magenta is S-. GLM for the average beta ztPRP between 0.5 and 2.5 sec found statistically significant differences for both CA1 and mPFC for both naïve vs. proficient, rewarded vs. unrewarded odorant and for the interaction between naïve vs. proficient and rewarded vs. unrewarded odorant (p<0.001, 376 observations, 368 d.f., F-statistic=110-104, p<0.001, 6 mice, 8 odor pairs, Extended Data Fig. 6). GLM for the average gamma ztPRP between 0.5 and 2.5 sec found statistically significant differences for both CA1 and mPFC for both naïve vs. proficient (p<0.001), rewarded vs. unrewarded odorant (p<0.001) and for the interactions between naïve vs. proficient and rewarded vs. unrewarded odorant (p<0.001), naïve vs. proficient and peak vs. trough (p<0.05) and rewarded vs. unrewarded odorant and peak vs. trough (p<0.001, 376 observations, 368 d.f., F-statistic=88.9-94.5, p<0.001, 6 mice, 8 odor pairs, Extended Data Fig. 6). Asterisks denote significant difference between rewarded and unrewarded odorants determined post hoc with t or ranksum tests (p<pFDR). F and G. i and ii. Time courses for the p value calculated with a ranksum test of the difference between rewarded and unrewarded trials for the ztPRP or the licks. The ranksum test was calculated in time bins of 33 msec. Shown are bounded lines of the average p value for ztPRP or licks (±CI) calculated per odorant per mouse. Fi beta CA1, Fii gamma CA1, Gi beta mPFC, Gii gamma mPFC. iii,iv Divergence times between rewarded and unrewarded ztPRP and licks calculated from the ranksum p value time courses (see methods). A GLM for the divergence times found statistically significant differences for both CA1 and mPFC between peak and licks and trough and licks (p<0.001, 133 observations, 130 d.f., F-statistic=11.6-21, p<0.001, 6 mice, 8 odor pairs, Extended Data Fig. 6). Asterisks denote significant differences determined post hoc with t or ranksum tests (p<pFDR).

In order to compare the divergence of lick behavior with tPRP we compared the time course for the decrease of p value estimating divergence between S+ and S- for licks vs. tPRP. Figures 6D and E show the time course for the z normalized peak and trough tPRP (ztPRP) for naïve and proficient animals for the different bandwidths for CA1 (Figures 6Di,ii and 6Ei,ii) and mPFC (Figures 6Diii,iv and 6Eiii,iv). For proficient mice ztPRP diverged between rewarded and unrewarded odorant shortly after odor onset.

Consistent with the results in Figure 3 a GLM analysis for the average beta ztPRP between 0.5 and 2.5 sec found statistically significant differences for both CA1 and mPFC for naïve vs. proficient, rewarded vs. unrewarded odorant and for the interaction between naïve vs. proficient and rewarded vs. unrewarded odorant (p<0.001, 376 observations, 368 d.f., F-statistic=110-104, p<0.001, 6 mice, 8 odor pairs, Extended Data Fig. 6). GLM for the average gamma ztPRP between 0.5 and 2.5 sec found statistically significant differences for both CA1 and mPFC for naïve vs. proficient (p<0.001), rewarded vs. unrewarded odorant (p<0.001) and for the interactions between naïve vs. proficient and rewarded vs. unrewarded odorant (p<0.001), naïve vs. proficient and peak vs. trough (p<0.05) and rewarded vs. unrewarded odorant and peak vs. trough (p<0.001, 376 observations, 368 d.f., F-statistic=88.9-94.5, p<0.001, 6 mice, 8 odor pairs, Extended Data Fig. 6).

To estimate the time for divergence of ztPRP between rewarded and unrewarded trials in proficient mice we calculated the p value for a two tailed t test for ztPRP for each mouse for each odorant pair and compared it to the time course for the ranksum p value for divergence of lick behavior. As shown for CA1 and mPFC in Figures 6Fi,ii and Gi,ii there was a sharp decline in the p values shortly after addition of the odorant and the decrease in p value took place at earlier times of ztPRP compared to licks. Divergence time was computed as the time point after odorant onset where the p value dropped below 0.005 for >= 1.2 sec after addition of the odorant. Figures 6 Fiii,iv and Giii,iv show the time for divergence for peak and trough ztPRP compared to lick behavior. A GLM found statistically significant differences for both CA1 and mPFC for divergence time between peak ztPRP and licks and trough ztPRP and licks (p<0.001, 133 observations, 130 d.f., F-statistic=11.6-21, p<0.001, 6 mice, 8 odor pairs, Extended Data Fig. 6).

Asterisks denote significant differences determined post hoc with t or ranksum tests (p<pFDR).

### The time for divergence between rewarded and unrewarded trials differs between pre- and post-lick-referenced tPRP

In order to understand the relationship between licks and the power of beta and gamma referenced to the peak and trough of theta (tPRP) we sorted the time of occurrence of theta oscillation peaks and troughs with respect to the time of onset of adjacent licks for proficient animals. Figures 7A and B show the probability density (PD) for peak (Figure 7A) and trough (Figure 7B) occurrence timed with respect to adjacent licks. Consistent with studies in mPFC (Amarante et al., 2017; Amarante and Laubach, 2021) the average peaks tend to occur near the lick while the trough probability density show a bimodal distribution with peaks before and after the lick. We then calculated beta and gamma ztPRP time courses for peaks and troughs that occur before and after the lick (pre- and post-lick-referenced ztPRP). We found that the time course for pre-lick- referenced tPRP (Figures 7C and E) tended to diverge less between rewarded and unrewarded odorants than the post-lick-referenced tPRP (Figure 7D and F) and that the divergence was sustained for post-lick-referenced ztPRPs and transient for pre-lick- referenced ztPRPs (Figures 7C-F). GLM for the average beta or gamma ztPRP between 0.5 and 2.5 sec found statistically significant differences for both CA1 and mPFC for naïve vs. proficient (naïve are not shown Figure 7), rewarded vs. unrewarded odorant and pre-lick vs post-lick (p<0.001, 752 observations, 736 d.f., F-statistic=91- 123, p<0.001, 6 mice, 8 odor pairs, Extended Data Fig. 7).

**Figure 7.**
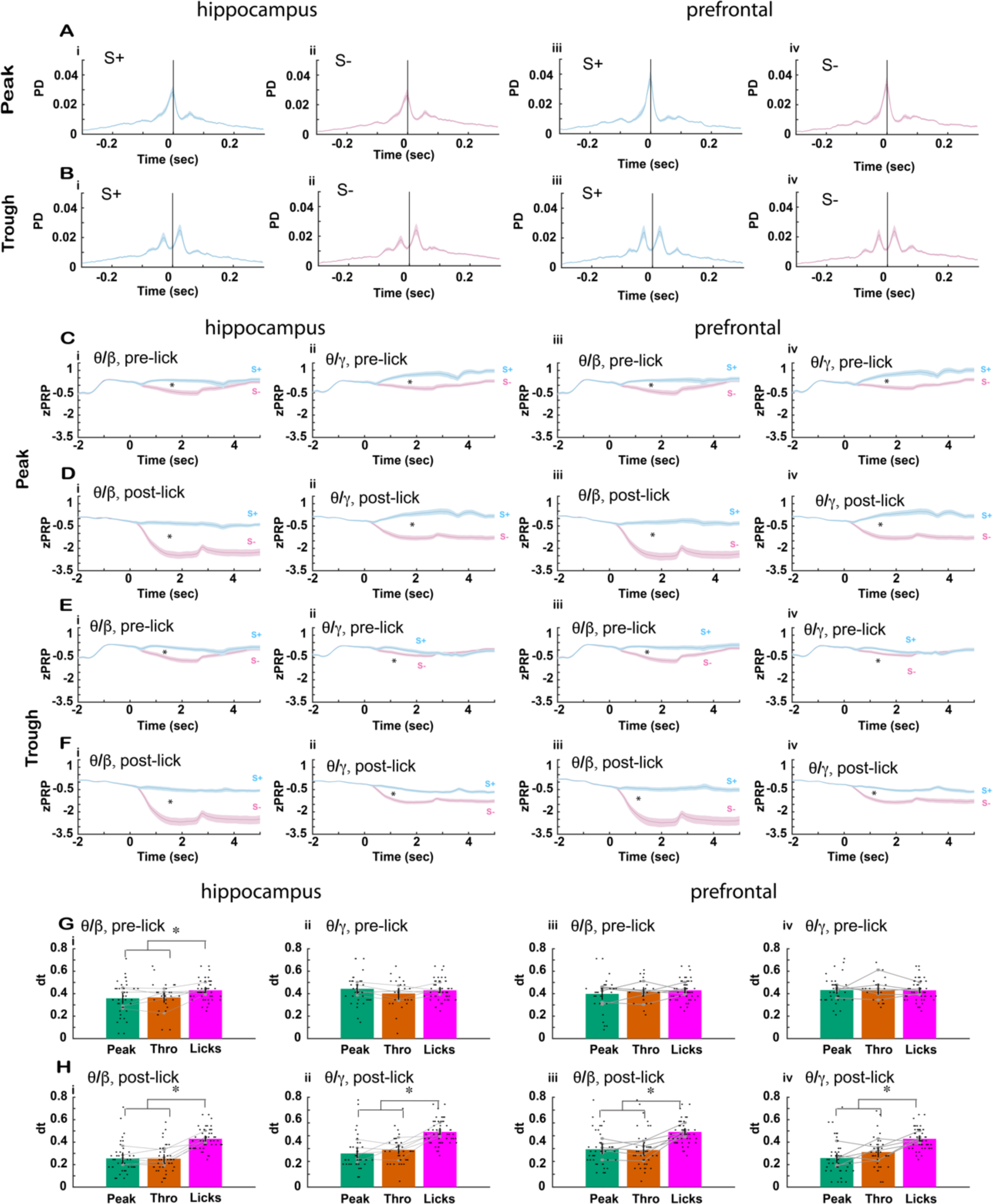
Lick-referenced tPRP for proficient mice. A to D. Plots of the probability density (PD) for the time of peaks (A) or troughs (B) of theta oscillations referenced to the time for adjacent lick onset shown for CA1 (i,ii) and mPFC (iii,iv) for S+ (Ai,Bi,Aiii,Biii) and S- (Aii,Bii,Aiv,Biv). The plots show a bounded line (<u>average+</u>CI) calculated from all the per mouse per odorant data. C-F. Time course for the average lick-referenced tPRP. C and D are for peak lick-referenced tPRP and E and F are for trough lick-referenced tPRP. C and E are pre-lick lick-referenced tPRP calculated for peaks and troughs that took place before the reference lick and D and F are post-lick lick-referenced tPRP calculated for peaks and troughs that took place after the reference lick. The tPRP is z normalized by subtracting the mean tPRPfrom -2 to 0 sec and dividing by the standard deviation in this period. The bounded line shows the average ztPRP bounded by the CI calculated from ztPRP traces per odorant per mouse (6 mice, 8 odor pairs). i, hippocampus theta/beta, ii, hippocampus theta/gamma, iii, mPFC theta/beta and iv, mPFC theta/gamma. Cyan is S+ and magenta is S-. GLM for the average beta or gamma ztPRP between 0.5 and 2.5 sec found statistically significant differences for both CA1 and mPFC for naïve vs. proficient (naïve are not shown in the figure), rewarded vs. unrewarded odorant and pre-lick vs post-lick (p<0.001, 752 observations, 736 d.f., F-statistic=91-123, p<0.001, 6 mice, 8 odor pairs, Extended Data Fig. 7). Asterisks denote significant difference between rewarded and unrewarded odorants determined post hoc with t or ranksum tests (p<pFDR). G and H. Divergence times between rewarded and unrewarded lick-referenced ztPRP and licks calculated from the p value time courses (see methods). G is pre-lick-referenced ztPRP and H is post-lick-referenced ztPRP. i and iii are for beta lick-referenced ztPRP and ii and iv are for gamma lick-referenced ztPRP. A GLM for the divergence times for pre-lick tPRP found no statistically significant differences between licks and either peak or trough for all bandwidths for mPFC (p>0.05, 100 observations, 97 d.f., F-statistic=0.006-0.65, p>0.05, 6 mice, 8 odor pairs, Extended Data Fig. 7). A GLM for the divergence times for pre-lick tPRP found no statistically significant differences between licks and either peak or trough for gamma for CA1 (p>0.05, 100 observations, 97 d.f., F- statistic=0.086, p>0.05, 6 mice, 8 odor pairs, Extended Data Fig. 7) and found a statistically significant difference between both peak and trough and licks for beta CA1 (p<0.05, 100 observations, 97 d.f., F-statistic=4, p<0.05, 6 mice, 8 odor pairs, Extended Data Fig. 7). In contrast, for post-lick ztPRP divergence for all bandwidths and for both CA1 and mPFC GLM found a statistically significant difference between both peak and trough and lick divergence (p<0.001, 124 observations, 121 d.f., F-statistic=17.8-31.8, p<0.001, 6 mice, 8 odor pairs, Extended Data Fig. 7). Asterisks denote significant differences determined post hoc with t or ranksum tests (p<pFDR).

Additionally, we found an interesting difference between pre-lick-referenced ztPRP and post-lick-referenced ztPRP when we assessed the time for divergence between rewarded and unrewarded trials for the time courses for lick-referenced ztPRP. For post-lick-referenced ztPRP the time for divergence was smaller than the time for divergence for lick behavior for all bandwidths for both peak and trough for both CA1 and mPFC (Figure 7H). In contrast, for pre-lick-referenced ztPRP in mPFC the time for divergence for both bandwidths and peak and trough did not differ from the time for divergence for lick behavior (Figure 7Giii,iv). For CA1 the time for divergence for pre-lick-referenced ztPRP for both peak and trough differed from the time for divergence for lick behavior for beta, but not for gamma (Figures 7Gi,ii). A GLM for the divergence times for pre-lick tPRP found no statistically significant differences between licks and either peak or trough for all bandwidths for mPFC (p>0.05, 100 observations, 97 d.f., F-statistic=0.006-0.65, p>0.05, 6 mice, 8 odor pairs, Extended Data Fig. 7). A GLM for the divergence times for pre-lick tPRP found no statistically significant differences between licks and either peak or trough for gamma for CA1 (p>0.05, 100 observations, 97 d.f., F-statistic=0.086, p>0.05, 6 mice, 8 odor pairs, Extended Data Fig. 7) and found a statistically significant difference between both peak and trough and licks for beta CA1 (p<0.05, 100 observations, 97 d.f., F-statistic=4, p<0.05, 6 mice, 8 odor pairs, Extended Data Fig. 7). In contrast, for post-lick ztPRP divergence for all bandwidths and for both CA1 and mPFC GLM found a statistically significant difference between both peak and trough and lick divergence (p<0.001, 124 observations, 121 d.f., F-statistic=17.8-31.8, p<0.001, 6 mice, 8 odor pairs, Extended Data Fig. 7).

### Coordinated hippocampal-prefrontal neural activity decreased for the unrewarded odorant as the animal became proficient

Coordinated hippocampal-prefrontal neural activity supports the organization of brain rhythms and is present during a range of cognitive functions presumably underlying transfer of information between these two brain regions (Colgin, 2011; Gordon, 2011; Headley and Paré, 2017; Lisman et al., 2017). We proceeded to determine whether there were changes in coordinated neural activity between dorsal CA1 and mPFC as the animal learned to discriminate the odorants. We used two complementary methods to quantify coordinated neural activity by calculating imaginary coherence (iCoherence) and phase locking value (PLV), measures of coherence that are independent of volume conduction (Bastos and Schoffelen, 2016; Namburi, 2021; Nolte et al., 2004).

Figure 8A shows an example of a spectrogram for the time course for iCoherence for a session where a mouse was engaged in the go-no go task. The pseudocolor plots show the average iCoherence time course per trial for (i) S+ naïve (ii) S- naïve (iii) S+ proficient (iv) S- proficient. For S+ proficient there is an increase in iCoherence after odorant addition compared to naïve. This positive iCoherence indicates that coherent oscillations that take place in CA1 before they ensue in mPFC. Another example for a different electrode pair from the same session for the proficient animal shows that the rewarded odorant elicits a decrease in theta iCoherence to a negative value (oscillations take place earlier in mPFC, Figure 8B). Figure 8C shows the histogram for odorant- elicited changes in theta iCoherence for all electrode pairs for the proficient mouse in this session. The distribution of the changes in iCoherence was broad and on the average the rewarded odorant elicited an increase in theta iCoherence and the unrewarded odorant elicited a smaller increase.

**Figure 8.**
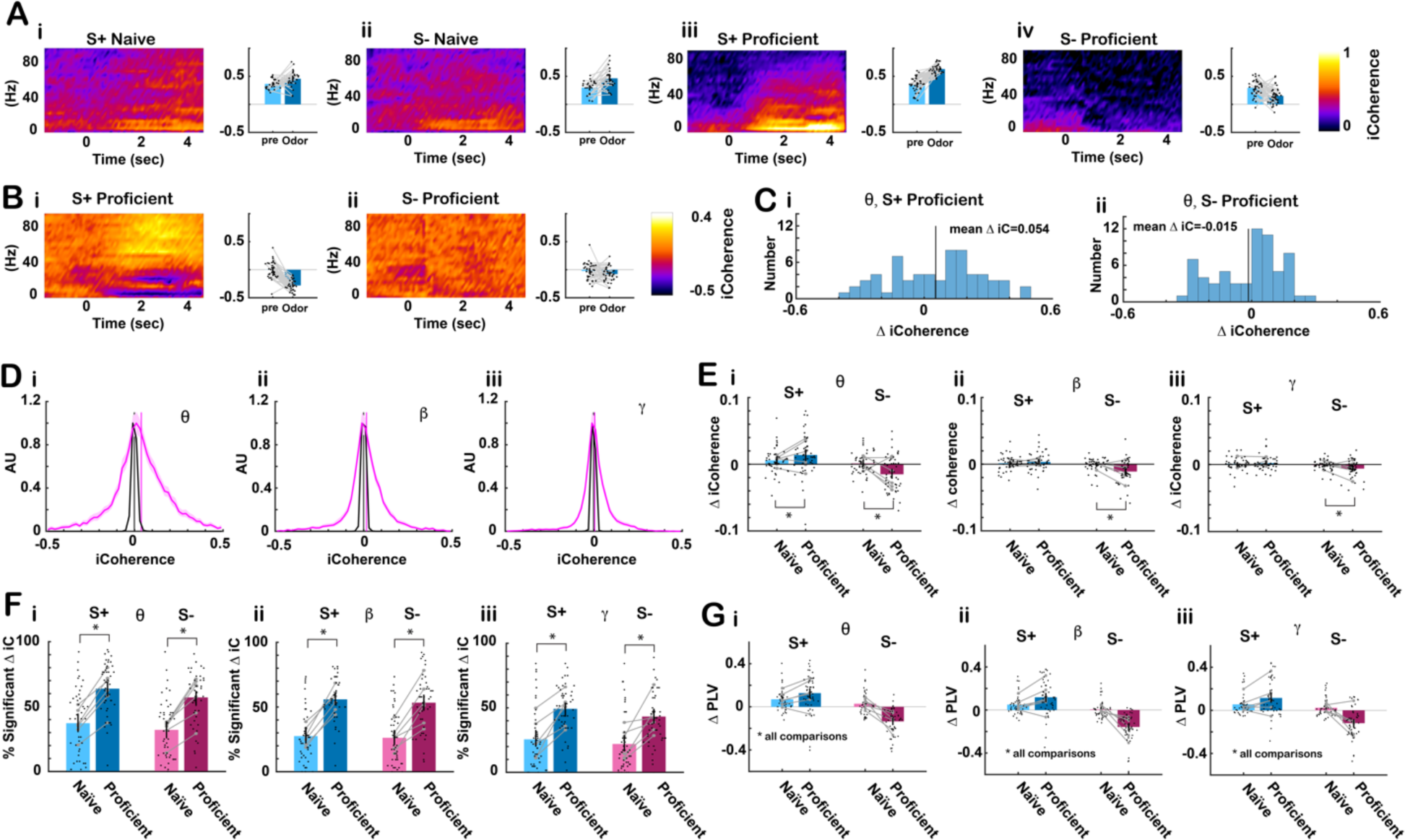
Odorant-elicited change in imaginary coherence and phase locking value decreased for the unrewarded odorant as the animal became proficient. A. Example of pseudocolor plots of the average mPFC/CA1 imaginary coherence (iCoherence) average trial time course for (i) S+ naïve (30 trials) (ii) S- naïve (27 trials) (iii) S+ proficient (33 trials) (iv) S- proficient (35 trials). Positive iCoherence means that CA1 oscillatory activity precedes the activity in mPFC because iCoherence is referenced to an electrode in the hippocampus. B. iCoherence time course for the same mPFC electrode referenced to a different CA1 electrode for the same proficient trials as in A. (i) S+ proficient (33 trials) (iv) S- proficient (35 trials). In this case mPFC precedes CA1. C. Odorant-elicited change in theta iCoherence for all 64 electrode pairs in the same session as in Aiii,iv. (i) S+ proficient (33 trials) (ii) S- proficient (35 trials). D. Distribution of average iCoherence during odorant application per mouse per odorant pair. The distribution is normalized to the peak. Pink is the distribution for iCoherence during the odorant and black is the distribution for iCoherence calculated after shuffling trials. Shown is the mean bounded by the 95% CI of the distributions calculated per odorant per mouse. All conditions were included in this distribution: naïve and proficient, S+ and S-. Vertical bars are the mean iCoherence values. E. Summary for changes in odorant-elicited changes in iCoherence (Δ iCoherence) as the animal learns. Average Δ iCoherence is shown for the different bandwidths (per mouse per odorant pair): (ii) theta, (iii) beta, (iv) gamma (6 mice, 8 odor pairs, the vertical line is the 95% CI). GLM found for all bandwidths a statistically significant difference for naïve vs. proficient (p<0.001, 188 observations, 184 d.f., F-statistic=6-12, p<0.001, 6 mice, 8 odor pairs, Extended Data Fig. 8) and for the interaction between naïve vs. proficient and rewarded vs. unrewarded (p<0.05, 188 observations, 184 d.f., F-statistic=6-12, p<0.001, 6 mice, 8 odor pairs, Extended Data Fig. 8). F. Percent significant odorant-elicited changes in iCoherence per odorant per mouse. GLM found for all bandwidths a statistically significant difference for naïve vs. proficient (p<0.001, 188 observations, 184 d.f., F-statistic=18.7-29.1, p<0.001, 6 mice, 8 odor pairs, Extended Data Fig. 8). G. Bar graphs showing odor-elicited change in average odorant-elicited change in phase- locking value (Δ PLV) per mouse per odor pair for (i) theta, (ii) beta, (iii) gamma. (6 mice, 8 odor pairs, the vertical line is the 95% CI). GLM found statistically significant differences between naïve vs. proficient (p<0.05) and for S+ vs. S- and the interaction between S+ vs. S- and naïve vs. proficient (p<0.001, 192 observations, 188 d.f., F- statistic=29.5-46.5, p<0.001, 6 mice, 8 odor pairs, Extended Data Fig. 8).

We proceeded to characterize iCoherence and the odorant-elicited changes in iCoherence for all sessions. The magenta bounded lines in Figure 8D show the distribution of average theta, beta and gamma iCoherence per mouse per odor pair during the odorant administration period (0.5 to 2.5 sec). The mean theta iCoherence illustrated by a vertical magenta line is positive, indicating that on the average theta oscillations take place earlier in CA1, consistent with findings by other investigators (Adhikari et al., 2010). Shuffling the trials results in a narrow symmetrical distribution for iCoherence centered at zero (Figure 8D, black bounded lines). Figure 8E shows the change of iCoherence (Δ iCoherence) elicited by the rewarded (S+) and unrewarded (S-) odorants for naïve and proficient mice calculated as the mean per odorant pair per mouse during the odorant administration period (0.5 to 2.5 sec). Δ iCoherence decreases for the unrewarded odorant when the animal becomes proficient. A GLM analysis found for all bandwidths statistically significant differences for Δ coherence for S+ vs. S- and the interaction between S+ vs. S- and naïve vs. proficient (p<0.001, 190 observations, 186 d.f., F-statistic=21.6-32.4, p<0.001, 6 mice, 8 odor pairs, Extended Data Fig. 8) and for theta the GLM analysis found additionally a significant difference between naïve and proficient (p<0.05).

Figure 8G shows the odor-elicited change in average PLV (Δ PLV) per mouse per odor pair during the odorant administration period (0.5 to 2.5 sec) for (i) theta, (ii) beta, (iii) gamma. Here we also found a negative Δ PLV for the unrewarded odorant for the proficient mice. For Δ PLV GLM found statistically significant differences for Δ PLV between naïve vs. proficient (p<0.05), S+ vs. S- and the interaction between S+ vs. S- and naïve vs. proficient (p<0.001, 192 observations, 188 d.f., F-statistic=29.5-46.5, p<0.001, 6 mice, 8 odor pairs, Extended Data Fig. 8). In conclusion, both the Δ iCoherence and Δ PLV measures indicate that as the mouse becomes proficient there was a decrease in coordinated hippocampal-prefrontal neural activity for the unrewarded odorant.

### Decreased performance for homozygote and heterozygote CaMKIIα knockout mice in the go-no go task

Since CaMKIIα is a protein involved in LTP we asked whether behavioral performance differed between the different CaMKIIα genotypes (WT, CaMKIIα Het and CaMKIIα KO). We trained mice from the three genotypes in the go-no go task. There was no difference in the number of sessions to criterion (Figure 9 A, GLM p>0.05, 24 observations, 21 d.f., F-statistic=0.26, p>0.05, 8 odor pairs, Extended Data Fig. 9). However, for proficient mice the percent correct was higher for WT compared to both CaMKIIα Het and CaMKIIα KO (Figure 9 B, GLM p<0.05, 137 observations, 134 d.f., F-statistic=3.7, p<0.05, 6 mice, 8 odor pairs, Extended Data Fig. 9) and the intertrial interval was larger for CaMKIIα Het (Figure 9 B, p<0.001, 137 observations, 134 d.f., F-statistic=13.5, p<0.001, 6 mice, 8 odor pairs, Extended Data Fig. 9).

**Figure 9.**
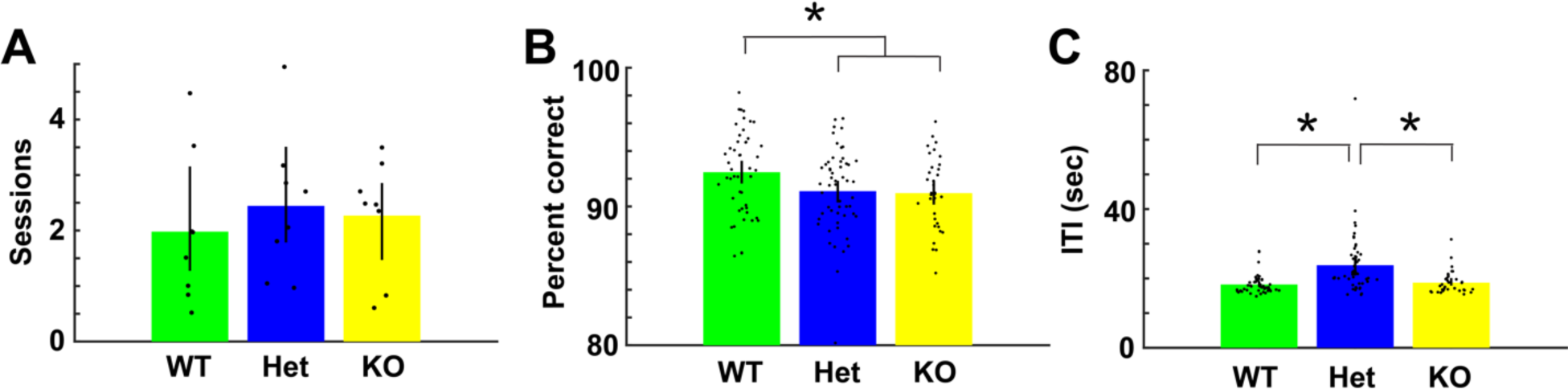
Analysis of the behavioral performance for mice of the three genotypes. (A). Number of sessions to criterion per odorant pair (2 blocks of 20 trials with performance >80%). There was no difference between genotypes in the number of sessions to criterion (GLM p>0.05, 24 observations, 21 d.f., F-statistic=0.26, p>0.05, 8 odor pairs). (B) Percent correct for proficient mice per odorant per mouse. GLM found a statistical difference for WT compared to both CaMKIIα Het and CaMKIIα KO (GLM p<0.05, 137 observations, 134 d.f., F-statistic=3.7, p<0.05, 6 mice, 8 odor pairs). (C). Intertrial interval (ITI) for proficient mice. GLM analysis found that the ITI for CaMKIIα Het differed from both CaMKIIα Het and CaMKIIα KO (p<0.001, 137 observations, 134 d.f., F-statistic=13.5, p<0.001, 6 mice, 8 odor pairs). Asterisks denote statistically significant differences evaluated with either t test or ranksum corrected for multiple comparisons (p<pFDR).

### The strength of tPAC and peak angle variance differed between the CaMKIIα genotypes

We then asked whether tPAC in the hippocampus and mPFC differs between the different genotypes. When we compared strength of tPAC, measured by the modulation index (MI), and the peak angle variance between genotypes (WT, CaMKIIα Het and CaMKIIα KO), we found significant differences. Figure 10 A, B shows the average MI per mouse per odorant pair for beta (Ai), high gamma (Bi) for hippocampus and for beta (Aii) and gamma (Bii) for prefrontal cortex. For beta tPAC GLM found a statistically significant difference for MI for the interaction between WT vs. CaMKIIα KO and S+ vs. S- in both the hippocampus and mPFC (p<0.001, 544 observations, 532 d.f., F- statistic=10.3-34.3, p<0.001, 6 mice, 8 odor pairs, Extended Data Fig. 10). For high gamma tPAC there was an increase in MI for CaMKIIα Het and a decrease in MI for CaMKIIα KO compared to WT and GLM analysis found a statistically significant difference between WT vs. CaMKIIα KO and WT vs. CaMKIIα Het for both hippocampus and mPFC (p<0.001, 544 observations, 532 d.f., F-statistic=10.3-34.3, p<0.001, 6 mice, 8 odor pairs, Extended Data Fig. 10).

**Figure 10.**
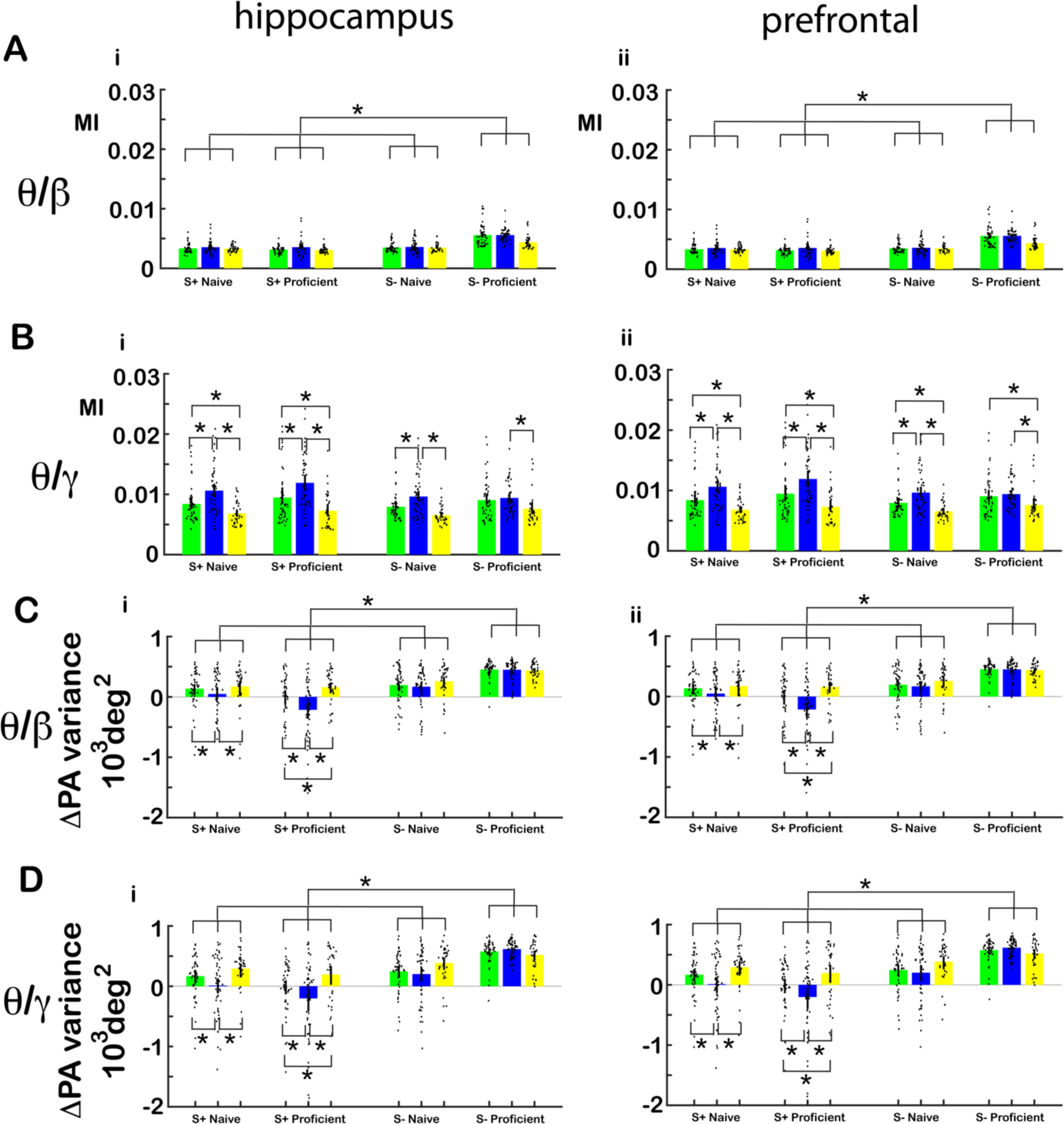
tPAC and peak angle variance per genotype. (A,B) Bar graphs showing average MI per mouse per odor pair for each genotype, S+ and S- in naïve and proficient mice for (A) theta/beta and (B) theta/gamma for (i) hippocampus and (ii) mPFC. The bars show the average MI, and the points are the MI per mouse per odor pair. For theta/beta tPAC MI GLM found a statistically significant difference for S+ vs. S-, for the interaction of WT vs. KO and S+ vs. S- and the interaction of WT vs. KO and naïve vs. proficient (p<0.001, 544 observations, 532 d.f., F-statistic=10.3-34.3, p<0.001, 6 mice, 8 odor pairs, Table S10). For theta/gamma tPAC MI GLM found a statistically significant difference between WT vs. KO and WT vs. Het (p<0.01) and for the interaction of WT vs. Het and S+ vs. S- (p<0.05, 544 observations, 532 d.f., F-statistic=10.3-34.3, p<0.001, 6 mice, 8 odor pairs, Table S10). (C,E) Bar graphs showing average peak angle variance per mouse per odor pair for each genotype, S+ and S- in naïve and proficient mice for (A) theta/beta and (B) theta/gamma for (i) hippocampus and (ii) mPFC. For beta tPAC MI GLM found a statistically significant difference for WT vs. Het (p<0.001) and WT vs. KO (p<0.05), S+ vs. S-, (p<0.001) and naïve vs. proficient (p<0.05, 544 observations, 532 d.f., F- statistic=20.6, p<0.001, 6 mice, 8 odor pairs, Table S11). For gamma tPAC MI GLM found a statistically significant difference for WT vs. Het (p<0.05) and WT vs. KO (p<0.05), S+ vs. S-, (p<0.001) and naïve vs. proficient (p<0.05, 544 observations, 532 d.f., F-statistic=18.8, p<0.001, 6 mice, 8 odor pairs, Table S11).

Figure 10 C, D shows average peak angle variance per mouse per odorant pair for beta (Ci), high gamma (Ei) in the hippocampus and (Cii) beta, (Eii) high gamma in prefrontal cortex. For proficient mice the peak angle variance for the rewarded odorant decreased for CaMKα Het and increased for CaMKα KO. GLM analysis found a statistically significant difference for WT vs. CaMKIIα Het, WT vs. CaMKIIα KO, naïve vs. proficient and S+ vs. S- for theta/beta and theta/gamma for both the hippocampus and mPFC (p<0.05, 544 observations, 532 d.f., F-statistic=20.6, p<0.001, 6 mice, 8 odor pairs, Extended Data Fig. 10).

### The accuracy for decoding the contextual identity of the odorant decreased in the CaMKIIα knockout mouse and was correlated with percent correct discrimination

The differences in tPAC between CaMKIIα genotypes raises the question whether decoding of contextual identity from tPRP is altered in the CaMKIIα KO and the CaMKIIα Het mice. Figure 11 shows the results of our comparison of decoding accuracy between genotypes. Figures 11A and B. show examples for proficient mice of the time course for the accuracy of decoding of odorant contextual identity by LDA trained using tPRP for the EAPA odor pair (Figure 11A theta/beta tPRP, Figure 11B theta/gamma tPRP, i. WT, ii. Het, iii. KO). For the WT and CaMKIIα Het mice the accuracy increases monotonously during the odorant epoch (Figures 11Ai,ii and 11Bi,ii) while for the CaMKIIα KO mouse the accuracy reaches a maximum value after one second and then decreases slightly for the rest of the odorant epoch (Figures 11Aiii and 11Biii). Figures 11C and D show the differences in decoding accuracy between the different genotypes assessed in the window from 1.5 to 2.5 sec after addition of the odorant. For both beta and gamma peak or trough tPAC decoding for both the hippocampus and the mPFC decoding accuracy was lowest for CaMKIIα KO. In addition, for gamma trough tPRP decoding for mPFC the accuracy was highest for WT, and decreased for both CaMKIIα Het and CaMKIIα KO mice (Figure 11Div). For beta tPRP LDA GLM found statistically significant differences for decoding accuracy between WT and KO for all conditions (p<0.001) and between WT and Het for peak gamma tPRP in the hippocampus, trough gamma tPRP in mPFC (p<0.05, 139 observations, 136 d.f., F-statistic=9.9-34.3, p<0.001, 6 mice, 8 odor pairs, Extended Data Fig. 11).

**Figure 11.**
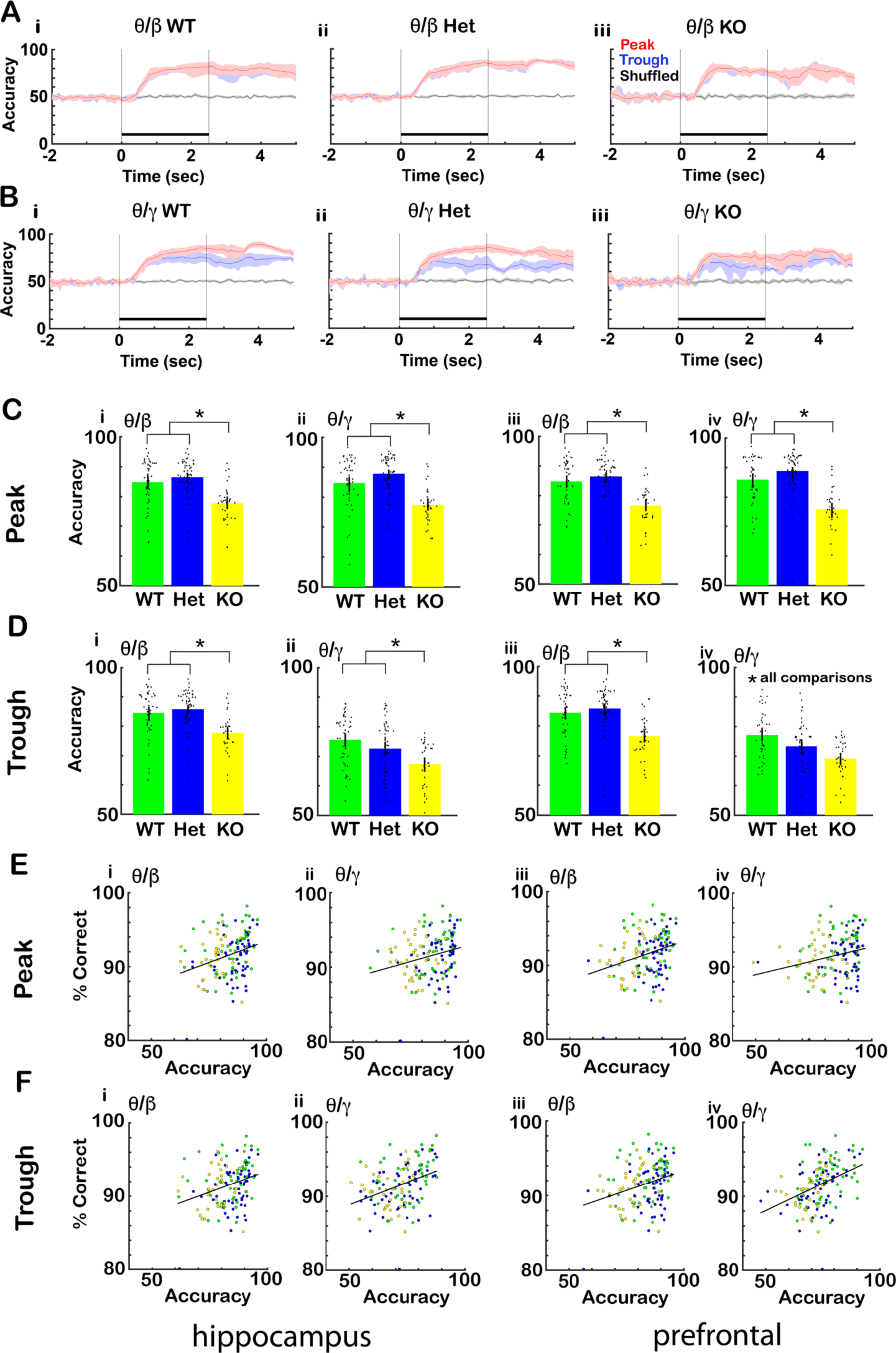
The accuracy for decoding the contextual identity of the odorant from tPRP decreased in the CaMKIIα knockout mouse and was correlated with percent correct discrimination. A and B. Examples for one mouse of the time course for the accuracy of odorant identification by linear discriminant analysis trained using CA1 tPRP for the EAPA odor pair. A. beta tPRP. B. gamma tPRP. (i) WT, (ii) Het, (iii) KO. Shadow: confidence interval, black bar: odorant application. C and D. Bar graphs showing the differences in discriminant accuracy between the different genotypes. C: Accuracy for peak tPRP for (i) theta/beta in the hippocampus, (ii) theta/gamma in the hippocampus, (iii) theta/beta in mPFC, (iv) theta/gamma in mPFC. D: Accuracy for through for (i) theta/beta in the hippocampus, (ii) theta/gamma in the hippocampus, (iii) theta/beta in mPFC, (iv) theta/gamma mPFC. The bars show the average accuracy, and the points are the accuracy per mouse per odor pair. The vertical bars show the confidence interval. For beta tPRP LDA GLM found statistically significant differences between WT and KO for all conditions (p<0.001) and between WT and Het for gamma trough tPRP (p<0.05, 756 observations, 744 d.f., F-statistic=9.9-34.3, p<0.001, 6 mice, 8 odor pairs, Supplementary Table 11). Asterisks show significant p values (p<pFDR) for post-hoc pairwise tests. E. Relationship for proficient mice between percent correct in the go-no go behavior and accuracy of odor identification by the LDA decoding algorithm shown per mouse per odor pair (6 mice, 8 odor pairs). The correlation coefficients were: E (i) 0.3, E (ii) 0.24, E (iii) 0.29, E (iv) 0.23, F (i) 0.3, F (ii) 0.36, F (iii) 0.29, F (iv) 0.40, and the p value for significance was p<0.01. Lines are best fit lines.

We then asked whether there was a relationship between contextual odorant identity decoding accuracy and percent correct performance for proficient mice in the go-no go task. Figures 11E and 11F show that there were statistically significant correlations between decoding accuracy and percent correct performance for all the different conditions. The correlation coefficients were as follows: 0.3 for hippocampal peak beta tPRP, 0.24 for hippocampal peak gamma tPRP, 0.29 for mPFC peak beta tPRP, 0.23 for mPFC peak gamma tPRP, 0.3 for hippocampal trough beta tPRP, 0.36 for hippocampal trough gamma tPRP, 0.29 for mPFC trough beta tPRP and 0.40 for mPFC trough gamma tPRP, and the p value for significance of the correlation coefficient was p<0.01. This indicates that the decoding accuracy obtained with tPRP is related to behavioral performance.

Finally, we did not find differences between CaMKIIα genotypes for divergence times between rewarded and unrewarded ztPRP or lick time courses (Figure 12, divergence times were calculated as in Figure 6). GLM for the divergence times did not find statistically significant differences between genotypes for licks (p>0.05, 127 observations, 124 d.f., F-statistic=1.5, p>0.05, 6 mice, 8 odor pairs, Extended Data Fig. 12). A GLM for the divergence times did not find statistically significant differences between genotypes for beta or gamma ztPRP (p>0.05, 252 observations, 246 d.f., F- statistic=0.5-1, p>0.05, 6 mice, 8 odor pairs, Extended Data Fig. 12).

**Figure 12.**
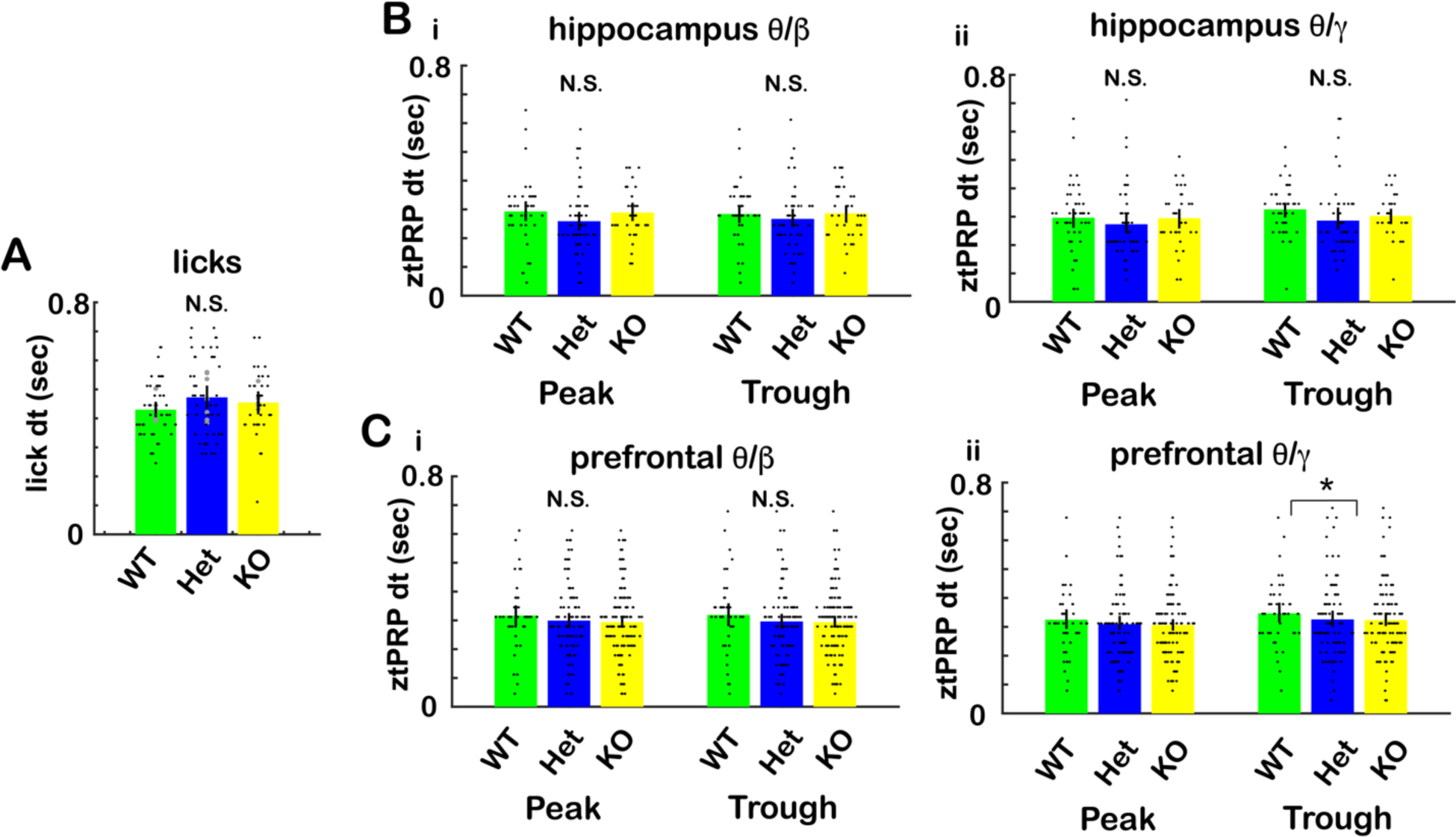
Divergence times between the ztPRP time courses for rewarded and unrewarded trials for the different CaMKIIα genotypes. Divergence times between rewarded and unrewarded ztPRP and licks were calculated from the ranksum p value time courses as shown in Figure 6 (also see methods). A. Divergence times for licks. A GLM for the divergence times did not find statistically significant differences between genotypes for licks (p>0.05, 127 observations, 124 d.f., F-statistic=1.5, p>0.05, 6 mice, 8 odor pairs, Extended Data Fig. 12). B and C Divergence times for ztPRP. B CA1, C. mPFC. i. beta tPRP, ii. gamma tPRP. A GLM for the divergence times did not find statistically significant differences between genotypes for beta or gamma ztPRP (p>0.05, 252 observations, 246 d.f., F-statistic=0.5-1, p>0.05, 6 mice, 8 odor pairs, Extended Data Fig. 12).

### Coherent hippocampal-prefrontal neural activity differed between CaMKIIα genotypes

We asked whether there were differences for coordinated hippocampal-prefrontal neural activity for the different CaMKIIα genotypes. Figure 13A shows the odorant-elicited changes in Δ iCoherence for the different genotypes. For theta Δ iCoherence GLM found a statistically significant difference between WT and KO (p<0.05, 544 observations, 532 d.f., F-statistic=5.35, p<0.001, 6 WT mice, 7 Het mice and 5 KO mice, 8 odor pairs, see Extended Data Fig. 13). However, these were relatively small differences of theta Δ iCoherence between genotypes and post-hoc tests did not yield significant differences between CaMKIIα KO and WT. Furthermore, the percent of electrode pairs that showed a significant Δ iCoherence was higher for Hets compared to WT for the rewarded odorant for beta and gamma Δ iCoherence (Figures 13Bii, iii). For beta Δ iCoherence GLM found a statistically significant difference between WT and Het (p<0.05, 544 observations, 532 d.f., F-statistic=18.1, p<0.001, 6 WT mice, 7 Het mice and 5 KO mice, 8 odor pairs, see Extended Data Fig. 13). For gamma Δ iCoherence GLM found a statistically significant difference for the interaction between WT and Het and rewarded vs. unrewarded odorant (p<0.05, 544 observations, 532 d.f., F-statistic=14.7, p<0.001, 6 WT mice, 7 Het mice and 5 KO mice, 8 odor pairs, see Extended Data Fig. 13).

**Figure 13.**
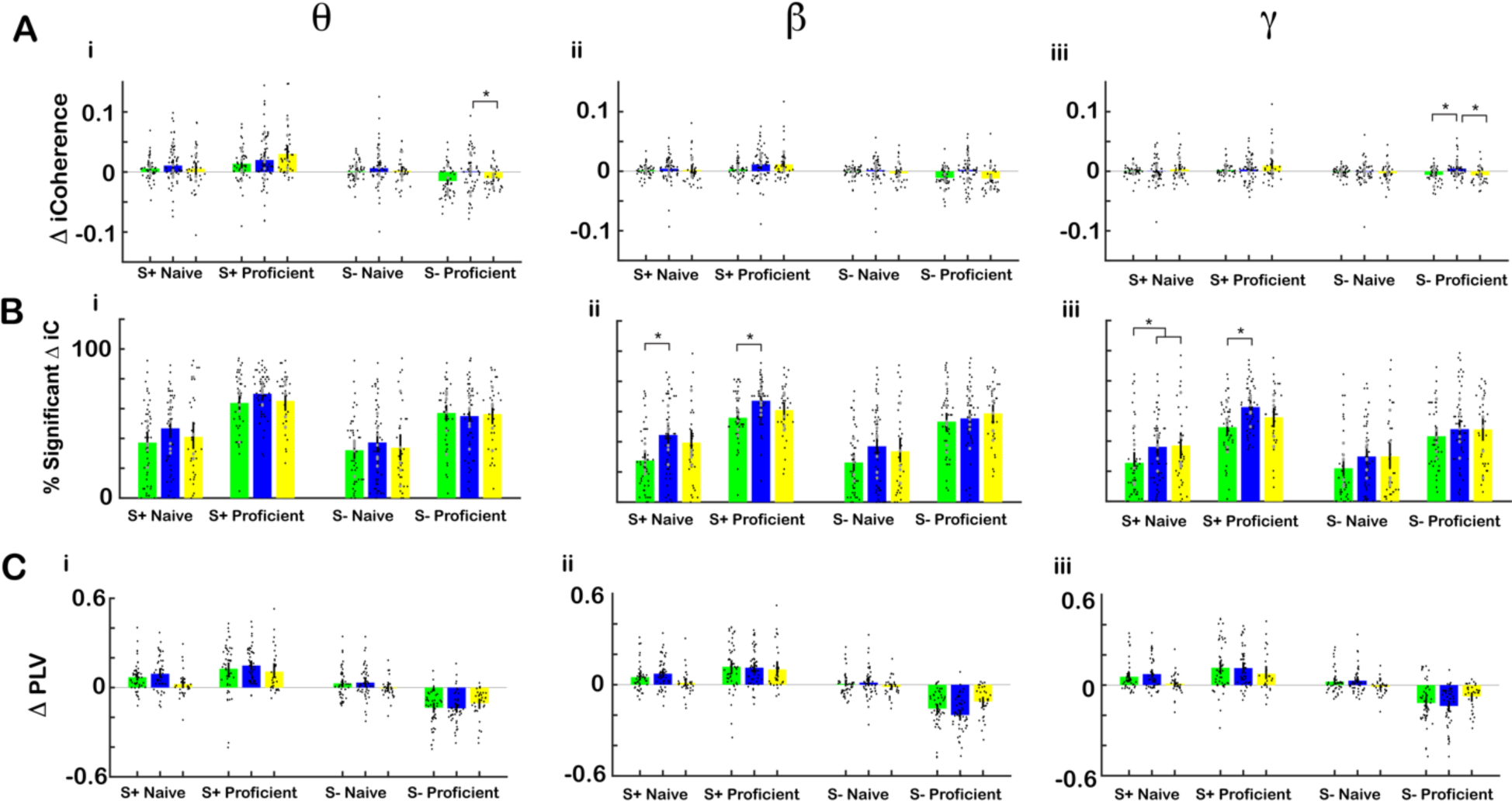
There were small differences in odorant-elicited change in imaginary coherence between CaMKIIα genotypes. A. Summary bar graphs comparing the odorant-elicited change in imaginary coherence (Δ iCoherence) for the different genotypes. (i) theta, (ii) beta, (iii) gamma. The bars show the average Δ iCoherence, and the points are Δ iCoherence per mouse per odor pair. The vertical bars show the confidence interval. For theta Δ iCoherence GLM found a statistically significant difference between WT and KO (p<0.05, 544 observations, 532 d.f., F-statistic=5.35, p<0.001, 6 WT mice, 7 Het mice and 5 KO mice, 8 odor pairs, see Extended Data Fig. 13). For gamma Δ iCoherence GLM found a statistically significant difference for the interaction between WT and KO and rewarded vs. unrewarded odorant (p<0.05, 544 observations, 532 d.f., F-statistic=2.76, p<0.05, 6 WT mice, 7 Het mice and 5 KO mice, 8 odor pairs, see Extended Data Fig. 12). Asterisks show significant p values (p<pFDR) for post-hoc pairwise tests. B. Percent significant odorant- elicited changes in iCoherence per odorant per mouse. For beta Δ iCoherence GLM found a statistically significant difference between WT and Het (p<0.05, 544 observations, 532 d.f., F-statistic=18.1, p<0.001, 6 WT mice, 7 Het mice and 5 KO mice, 8 odor pairs, see Extended Data Fig. 13). For gamma Δ iCoherence GLM found a statistically significant difference for the interaction between WT and het and rewarded vs. unrewarded odorant (p<0.05, 544 observations, 532 d.f., F-statistic=14.7, p<0.001, 6 WT mice, 7 Het mice and 5 KO mice, 8 odor pairs, see Extended Data Fig. 12). C. Summary bar graphs comparing change in Δ PLV for the different genotypes. (i) beta, (ii) theta, (iii) gamma. For Δ PLV GLM found no statistically significant differences between WT and KO, and for the interaction of WT vs. KO and S+ vs. S- (p>0.05, 512 observations, 500 d.f., F-statistic=25.5-39.9, p<0.001, 6 mice, 8 odor pairs, see Extended Data Fig. 13).

Δ PLV did not show any differences between genotypes (Figure 13C). The GLM found no statistically significant differences for Δ PLV between WT and KO or WT and Het (p>0.05, 512 observations, 500 d.f., F-statistic=25.5-39.9, p<0.001, 6 mice, 8 odor pairs, Extended Data Fig. 13).

### Acute block of CaMKIIα/β by local infusion of KN93 into CA1 elicits an increase in the accuracy for decoding the contextual identity of the odorant from tPRP

We performed go-no go experiments in mice that were perfused bilaterally in dorsal CA1 with either the CaMKIIα/β inhibitor KN93 (10 μM, 200 nl per site) or with the chemical analogue KN92 that does not inhibit CaMKIIα/β (Barcomb et al., 2015; Burgdorf et al., 2017). We decoded the contextual identity of the odorant in proficient mice from tPRP using LDA with the same procedure as in Figure 11. Figure 14A shows that KN93 elicited an increase in the accuracy of decoding the contextual identity of the odorant for the odorant application period (0.5-2.5 sec). GLM found statistically significant differences for accuracy between KN93 and KN92 (control) for beta and gamma tPRP (p<0.001, 26 observations, 22 d.f., F-statistic=7-9, p<0.05, 6 mice, 2 odor pairs, Extended Data Fig. 14). Post-hoc analysis did not find significant differences. In addition, Figures 14B shows that for beta tPRP the time for discrimination of the difference between ztPRP in rewarded and unrewarded trials was faster for KN93 than KN92 (control). Discrimination time was determined using the same method as in Figure GLM found statistically significant differences for accuracy between KN93 and KN92 (control) for beta tPRP (p<0.05, 18 observations, 15 d.f., F-statistic=4.6, p<0.05, 6 mice, 2 odor pairs, Extended Data Fig. 14). Post-hoc analysis did not find significant differences. These data indicate that acute inhibition of CaMKIIα/β in dorsal CA1 yields different results for decoding of contextual odorant identity and discrimination time compared to CaMKIIα KO and Hets (compare Figure 14 with Figure 11 for decoding and with Figure 12 for discrimination time).

**Figure 14.**
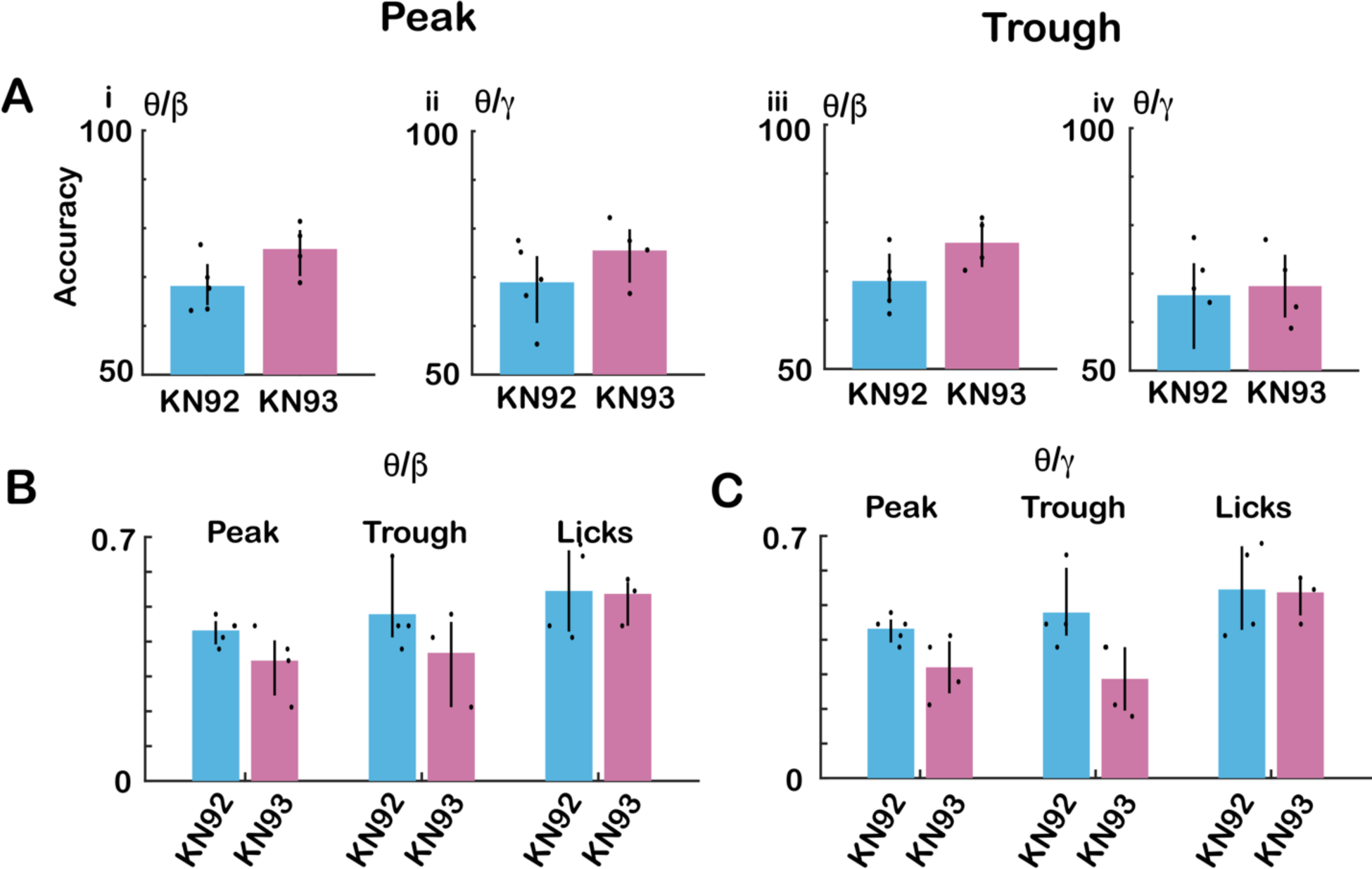
Accuracy for decoding of contextual identity and divergence time between rewarded and unrewarded trials for ztPRP upon acute block of CaMKIIα/β by local infusion of KN93 into CA1. A. Accuracy for decoding of contextual identity of the odorant for the odorant period (0.5 to 2.5 sec) determined by LDA analysis of tPRP (as performed in Figure 11) for mice perfused with KN93 or KN92 (control). i. peak beta tPRP, ii. peak gamma tPRP, iii trough beta tPRP and iv. trough gamma tPRP. GLM found statistically significant differences for accuracy between KN93 and KN92 (control) for beta and gamma tPRP (p<0.001, 26 observations, 22 d.f., F-statistic=7-9, p<0.05, 6 mice, 2 odor pairs, Extended Data Fig. 14). Post-hoc analysis did not find significant differences. B and C. Time for divergence for the time course of ztPRP between rewarded and unrewarded trials, determined as in Figure 6, for mice perfused with KN93 or KN92 (control). B beta ztPRP, C. gamma ztPRP. GLM found statistically significant differences for accuracy between KN93 and KN92 (control) for beta tPRP (p<0.05, 18 observations, 15 d.f., F-statistic=4.6, p<0.05, 6 mice, 2 odor pairs, Extended Data Fig. 14). There was no statistically significant difference for gamma tPRP (p>0.05). Post-hoc analysis did not find significant differences.

## Discussion

Santiago Ramon y Cajal described the hippocampus as a quaternary region of the olfactory system (Cajal, 1904), but subsequent studies showed that the hippocampus is involved in learning and memory in non-olfactory tasks (Nakazawa et al., 2004).

However, the hippocampus does play a role in olfactory learning. Indeed, calbindin-expressing pyramidal cells in dorsal CA1 develop selective responses to odorants as the animal becomes proficient in the go-no go olfactory discrimination task (Li et al., 2017). Furthermore, studies of oscillatory neural activity implicate directional coupling from the OB to the hippocampus in go-no go learning (Gourevitch et al., 2010; Martin et al., 2007). In addition, Granger directionality analysis found that distinct low frequency oscillation bandwidths link the OB and hippocampus (Nguyen Chi et al., 2016).

Interestingly, mPFC has also been proposed as a downstream brain area coupled with the OB. Coupling of low frequency oscillations between the OB and mPFC increases during freezing periods in auditory conditioned fear learning (Moberly et al., 2018), mPFC neurons represent odor value (Wang et al., 2020) and at rest there is strong coupling of low frequency oscillations for OB-hippocampus and OB-mPFC (Mofleh and Kocsis, 2021). Furthermore, beta and theta synchrony between mPFC and olfactory regions was elevated as rats switched their attention to odors (Cansler et al., 2021).

Taken together with the fact that we showed that when mice become proficient in discriminating odorants in the go-no go task contextual odorant identity can be decoded from beta and gamma tPRP (Losacco et al., 2020) these findings raise the question whether coupled oscillations in mPFC and the hippocampus play a role in olfactory discrimination in go-no go associative learning.

In this study we found that as animals became proficient in the go-no go task there was an increase in the variance of the peak angle for beta and high gamma tPAC for the unrewarded odorant (Figure 2) accompanied by a decrease in tPRP for this odorant (Figures 3 and 6) in CA1 and mPFC. This decrease in tPRP for S- was accompanied by a small increase in tPRP for S+ resulting in a sharp increase in accuracy for decoding of the contextual odorant identity for the proficient mouse (Figure 5). Furthermore, divergence in tPRP between rewarded and unrewarded trials took place before divergence in lick behavior (Figures 6 and 7). When we tested CaMKIIα KO mice we found a decrease in the accuracy of decoding of contextual odorant identity (Figure 11). Finally, the behavioral performance for CaMKIIα KO and CaMKIIα Het mice was lower than the performance of WT mice (Figure 9) and the accuracy for decoding of contextual odorant identity from tPAC correlated with behavioral performance (Figure 11). Odor-elicited changes in imaginary coherence decreased for the unrewarded odorant as the animal became proficient (Figure 8). These findings are consistent with a role for coordinated oscillatory neuronal activity in the hippocampal-mPFC axis in the go-no go olfactory discrimination task.

Theta frequency stimulation, eliciting intrinsic gamma frequency oscillations, is known to be essential for LTP (Bliss and Lomo, 1973; Butler et al., 2016; Larson and Munkácsy, 2015) and tPAC of higher frequency bursts is thought to be important for information transfer between brain regions which is thought to be essential for learning (Fries, 2005). In the hippocampus high frequency oscillations at different phases of theta carry different information. Indeed, Siegle and Wilson (2014) showed that mice increased performance in the encoding epoch for a spatial navigation task when parvalbumin interneurons were stimulated at the peak of theta whereas when these interneurons were stimulated at the trough of theta mice increased performance in the retrieval epoch. Interestingly, here we find that as the animal becomes proficient the variance of the peak angle of tPAC increases substantially for the unrewarded odorant (Figure 2). As a result, if the gamma or beta frequencies were being read by a downstream observer at a fixed angle, the information conveyed by oscillations elicited by the rewarded odorant would be missed because of the constantly changing theta phase angle. In contrast, the peak angle variance for the rewarded odorant is small, and presumably this would result in more faithful reading of this information. This is likely what underlies the increased accuracy for decoding the contextual odorant identity from beta and gamma tPRP in the proficient mouse (Figure 5). Importantly, respiration-coupled oscillations aid the exchange of information between OB and the hippocampus (Nguyen Chi et al., 2016). The high frequency (6-14 Hz) CA1 theta oscillations studied here may be caused by theta oscillatory input from the OB entrained by high frequency sniffing of the proficient animal in the go-no go task (Li et al., 2015), or may be due to complex interactions of the olfacto-hippocampal circuit. Future studies are needed to determine the precise role of sniffing vs. olfacto-hippocampal interactions in setting theta oscillations in CA1 during the go-no go task.

Full genetic knockout of CaMKIIα results in impairments in hippocampus-dependent cognitive tasks and impaired LTP and LTD in adult mice (Coultrap et al., 2014; Silva et al., 1992b). In contrast, although CaMKIIα Het mice share some behavioral deficits with the full CaMKIIα KO including deficient learning and memory in the Morris water maze (Frankland et al., 2001; Silva et al., 1996), adult CaMKIIα Het mice have additional working memory deficits such as repeat entry errors in the radial arms version of the Morris water maze (Matsuo et al., 2009; Yamasaki et al., 2008). In addition, while young (p 12-16) CaMKIIα Het show LTP of the same magnitude as WT mice in CA1, young adult mice display impaired basal synaptic transmission, but do not have a deficit in LTP (Goodell et al., 2016) suggesting the development of compensatory mechanisms for LTP in the Het. tPAC plays a role in hippocampal learning and memory (Siegle and Wilson, 2014) raising the question whether the CaMKIIα Het and CaMKIIα KO have deficits in cross frequency coupling.

Here we performed to our knowledge the first study to determine whether CaMKIIα- deficient mice display altered tPAC. We find that the accuracy for decoding of contextual odorant identity from beta and gamma tPRP is decreased in CaMKIIα KO mice, but not in CaMKIIα Hets (with the exception of a decrease accuracy for mPFC gamma trough tPRP decoding, Figure 11). Interestingly, the strength of tPAC, measured as the MI, was highest for gamma tPAC CaMKIIα Hets compared to both WT and CaMKIIα KO (Figure 10B). For CaMKIIα Hets the peak angle variance decreased significantly for the rewarded odorant when the animal became proficient (Figure 10C,D). If a downstream neural observer is evaluating contextual odorant identity by observing beta or gamma frequency bursts in phase with theta oscillations these changes in tPAC for CaMKIIα Hets would tend to increase the ability to discriminate (albeit with smaller accuracy for high gamma trough). Thus, the differences found for tPAC for CaMKIIα Hets may be compensatory leading to no difference in decoding accuracy between WT and CaMKIIα Hets (Figure 11). This would agree with the interpretation by Goodell et al. who indicated that LTP deficits in CaMKIIα Hets are restored by compensatory changes during development (Goodell et al., 2016). Finally, given that CaMKIIα Hets have a phenotype reminiscent of schizophrenia it is interesting that resting-state gamma tPAC has been found to be increased in patients with schizophrenia (Won et al., 2018). These authors speculate that increased tPAC may be related to the compensatory hyper-arousal patterns of the dysfunctional default-mode network in schizophrenia.

Whether the tPAC/tPRP changes found in CaMKIIα Het and CaMKIIα KO mice are due to the decreased expression of CaMKIIα protein or to developmental changes in circuits known to change in these mice such as in the dentate gyrus is an open question that will require future studies with temporally and spatially restricted changes in CaMKIIα activity. The result of inhibition of CaMKIIα/β (Figure 14) suggest that the effect of genotype is not exclusively due to inhibition of CaMKIIα/β in CA1. CaMKIIα may alter plasticity of pyramidal neurons in brain regions other than CA1. Furthermore, a decrease in CaMKIIα expression may alter the postsynaptic regulation by CaMKIIα of inhibitory synapse transmission onto the pyramidal neurons (Cook et al., 2021; Udakis et al., 2020). Finally, a subpopulation of granule cells that play a role in odorant discrimination in the go/no go task and are involved in generating the gamma frequency oscillatory activity in the bulb express CaMKIIα (Malvaut et al., 2017). The OB granule cells may be involved in changing coordinated oscillations between the OB and hippocampus in the CaMKIIα KO/Het mice.

In conclusion we found that as the mouse learns to differentiate odorants in the go-no go associative learning task there are changes in tPAC that result in an increase of the accuracy of decoding of the contextual odorant identity from tPRP. Finally, the accuracy of decoding the contextual odorant identity from tPRP decreased in the CaMKIIα KO, but did not decrease in the CaMKIIα Het and this decoding accuracy correlated with behavioral performance across genotypes.

## Acknowledgements

We thank Ms. Nicole Arevalo for animal husbandry and Ms. Dnate’ Baxter for laboratory support. We thank Dr. Stephen Coultrap for helpful discussion and animal support and Mrs. Lauraine Mediavillo for assistance in collecting data. This research was supported by an Administrative Supplement S1 to NIH UF1 NS116241 (to DRG), NIH R01 NS081248 (to KUB), NIH R01 DC000566 (to DR) and by a pilot grant from the Center for NeuroScience (CNS) of the University of Colorado School of Medicine (to KUB and DR).

## Extended data legends

Extended Data Fig. 2. Statistics for Figure 2.

Extended Data Fig. 3. Statistics for Figure 3.

Extended Data Fig. 4. Statistics for Figure 4.

Extended Data Fig. 5. Statistics for Figure 5.

Extended Data Fig. 6. Statistics for Figure 6.

Extended Data Fig. 7. Statistics for Figure 7.

Extended Data Fig. 8. Statistics for Figure 8.

Extended Data Fig. 9. Statistics for Figure 9.

Extended Data Fig. 10. Statistics for Figure 10.

Extended Data Fig. 11. Statistics for Figure 11.

Extended Data Fig. 12. Statistics for Figure 12.

Extended Data Fig. 13. Statistics for Figure 13.

Extended Data Fig. 14. Statistics for Figure 14

## References

Adhikari, A., Topiwala, M.A., and Gordon, J.A. (2010). Synchronized Activity between the Ventral Hippocampus and the Medial Prefrontal Cortex during Anxiety. Neuron 65, 257–269.

Agresti, A. (2015). Foundations of linear and generalized linear models. In Wiley series in probability and statistics (Hoboken, New Jersey: John Wiley & Sons Inc.,), pp. 1 online resource.

Amarante, L.M., Caetano, M.S., and Laubach, M. (2017). Medial Frontal Theta Is Entrained to Rewarded Actions. J Neurosci 37, 10757–10769.

Amarante, L.M., and Laubach, M. (2021). Coherent theta activity in the medial and orbital frontal cortices encodes reward value. eLife 10, e63372.

Barcomb, K., Goodell, D.J., Arnold, D.B., and Bayer, K.U. (2015). Live imaging of endogenous Ca(2)(+)/calmodulin-dependent protein kinase II in neurons reveals that ischemia-related aggregation does not require kinase activity. J Neurochem 135, 666–673.

Bastos, A.M., and Schoffelen, J.-M. (2016). A Tutorial Review of Functional Connectivity Analysis Methods and Their Interpretational Pitfalls. Frontiers in Systems Neuroscience 9.

Bayer, K.U., and Schulman, H. (2019). CaM Kinase: Still Inspiring at 40. Neuron 103, 380–394.

Bear, M.F., Cooke, S.F., Giese, K.P., Kaang, B.K., Kennedy, M.B., Kim, J.I., Morris, R.G.M., and Park, P. (2018). In memoriam: John Lisman - commentaries on CaMKII as a memory molecule. Mol Brain 11, 76.

Belluscio, M.A., Mizuseki, K., Schmidt, R., Kempter, R., and Buzsáki, G. (2012). Cross- Frequency Phase–Phase Coupling between Theta and Gamma Oscillations in the Hippocampus. The Journal of Neuroscience 32, 423.

Bliss, T.V., and Lomo, T. (1973). Long-lasting potentiation of synaptic transmission in the dentate area of the anaesthetized rabbit following stimulation of the perforant path. J Physiol 232, 331–356.

Borjigin, J., Lee, U., Liu, T., Pal, D., Huff, S., Klarr, D., Sloboda, J., Hernandez, J., Wang, M.M., and Mashour, G.A. (2013). Surge of neurophysiological coherence and connectivity in the dying brain. Proceedings of the National Academy of Sciences 110, 14432.

Buonviso, N., Amat, C., Litaudon, P., Roux, S., Royet, J.P., Farget, V., and Sicard, G. (2003). Rhythm sequence through the olfactory bulb layers during the time window of a respiratory cycle. EurJNeurosci 17, 1811–1819.

Burgdorf, C.E., Schierberl, K.C., Lee, A.S., Fischer, D.K., Van Kempen, T.A., Mudragel, V., Huganir, R.L., Milner, T.A., Glass, M.J., and Rajadhyaksha, A.M. (2017). Extinction of Contextual Cocaine Memories Requires Ca v1.2 within D1R-Expressing Cells and Recruits Hippocampal Ca v1.2-Dependent Signaling Mechanisms. The Journal of Neuroscience 37, 11894.

Butler, J.L., Mendonça, P.R.F., Robinson, H.P.C., and Paulsen, O. (2016). Intrinsic Cornu Ammonis Area 1 Theta-Nested Gamma Oscillations Induced by Optogenetic Theta Frequency Stimulation. The Journal of Neuroscience 36, 4155.

Cajal, S. (1904). Textura del sistema nervioso del hombre y de los vertebrados (Madrid: Moya).

Cansler, H.L., in ’t Zandt, E.E., Carlson, K.S., Khan, W.T., Ma, M., and Wesson, D.W. (2021). Organization and engagement of a prefrontal-olfactory network during olfactory selective attention. bioRxiv, 2021.2009.2004.458996.

Chen, C., Rainnie, D.G., Greene, R.W., and Tonegawa, S. (1994). Abnormal fear response and aggressive behavior in mutant mice deficient for alpha-calcium- calmodulin kinase II. Science 266, 291–294.

Chia, P.H., Zhong, F.L., Niwa, S., Bonnard, C., Utami, K.H., Zeng, R., Lee, H., Eskin, A., Nelson, S.F., Xie, W.H., et al. (2018). A homozygous loss-of-function CAMK2A mutation causes growth delay, frequent seizures and severe intellectual disability. eLife 7, e32451.

Colgin, L.L. (2011). Oscillations and hippocampal–prefrontal synchrony. Current Opinion in Neurobiology 21, 467–474.

Colgin, L.L. (2015). Theta-gamma coupling in the entorhinal-hippocampal system. Curr Opin Neurobiol 31, 45–50.

Colgin, L.L., and Moser, E.I. (2010). Gamma Oscillations in the Hippocampus. Physiology 25, 319–329.

Cook, S.G., Buonarati, O.R., Coultrap, S.J., and Bayer, K.U. (2021). CaMKII holoenzyme mechanisms that govern the LTP versus LTD decision. Sci Adv 7.

Coultrap, S.J., Freund, R.K., O’Leary, H., Sanderson, J.L., Roche, K.W., Dell’Acqua, M.L., and Bayer, K.U. (2014). Autonomous CaMKII mediates both LTP and LTD using a mechanism for differential substrate site selection. Cell Rep 6, 431–437.

Curran-Everett, D. (2000). Multiple comparisons: philosophies and illustrations. AmJPhysiol RegulIntegrComp Physiol 279, R1–R8.

Eleore, L., Lopez-Ramos, J.C., Guerra-Narbona, R., and Delgado-Garcia, J.M. (2011). Role of reuniens nucleus projections to the medial prefrontal cortex and to the hippocampal pyramidal CA1 area in associative learning. PLoS One 6, e23538.

Frankland, P.W., O’Brien, C., Ohno, M., Kirkwood, A., and Silva, A.J. (2001). Alpha- CaMKII-dependent plasticity in the cortex is required for permanent memory. Nature 411, 309–313.

Fries, P. (2005). A mechanism for cognitive dynamics: neuronal communication through neuronal coherence. Trends Cogn Sci 9, 474–480.

Fromer, M., Pocklington, A.J., Kavanagh, D.H., Williams, H.J., Dwyer, S., Gormley, P., Georgieva, L., Rees, E., Palta, P., Ruderfer, D.M., et al. (2014). De novo mutations in schizophrenia implicate synaptic networks. Nature 506, 179–184.

Goodell, D.J., Benke, T.A., and Bayer, K.U. (2016). Developmental restoration of LTP deficits in heterozygous CaMKIIα KO mice. Journal of Neurophysiology 116, 2140–2151.

Gordon, J.A. (2011). Oscillations and hippocampal–prefrontal synchrony. Current Opinion in Neurobiology 21, 486–491.

Gourevitch, B., Kay, L.M., and Martin, C. (2010). Directional coupling from the olfactory bulb to the hippocampus during a go/no-go odor discrimination task. J Neurophysiol 103, 2633–2641.

Halsey, L.G., Curran-Everett, D., Vowler, S.L., and Drummond, G.B. (2015). The fickle P value generates irreproducible results. Nature methods 12, 179–185.

Hasegawa, S., Furuichi, T., Yoshida, T., Endoh, K., Kato, K., Sado, M., Maeda, R., Kitamoto, A., Miyao, T., Suzuki, R., et al. (2009). Transgenic up-regulation of alpha- CaMKII in forebrain leads to increased anxiety-like behaviors and aggression. Mol Brain 2, 6.

Headley, D.B., and Paré, D. (2017). Common oscillatory mechanisms across multiple memory systems. npj Science of Learning 2, 1.

Kaplan, R., Bush, D., Bonnefond, M., Bandettini, P.A., Barnes, G.R., Doeller, C.F., and Burgess, N. (2014). Medial prefrontal theta phase coupling during spatial memory retrieval. Hippocampus 24, 656–665.

Klein, S.B., Cosmides, L., Tooby, J., and Chance, S. (2002). Decisions and the evolution of memory: multiple systems, multiple functions. Psychol Rev 109, 306–329.

Lachaux, J.P., Rodriguez, E., Martinerie, J., and Varela, F.J. (1999). Measuring phase synchrony in brain signals. Hum Brain Mapp 8, 194–208.

Larson, J., and Munkácsy, E. (2015). Theta-burst LTP. Brain Research 1621, 38–50.

Li, A., Gire, D.H., and Restrepo, D. (2015). Υ spike-field coherence in a population of olfactory bulb neurons differentiates between odors irrespective of associated outcome. J Neurosci 35, 5808–5822.

Li, Y., Xu, J., Liu, Y., Zhu, J., Liu, N., Zeng, W., Huang, N., Rasch, M.J., Jiang, H., Gu, X., et al. (2017). A distinct entorhinal cortex to hippocampal CA1 direct circuit for olfactory associative learning. Nature Neuroscience 20, 559–570.

Lisman, J., Buzsaki, G., Eichenbaum, H., Nadel, L., Ranganath, C., and Redish, A.D. (2017). Viewpoints: how the hippocampus contributes to memory, navigation and cognition. Nat Neurosci 20, 1434–1447.

Lisman, J., Yasuda, R., and Raghavachari, S. (2012). Mechanisms of CaMKII action in long-term potentiation. Nat Rev Neurosci 13, 169–182.

Losacco, J., Ramirez-Gordillo, D., Gilmer, J., and Restrepo, D. (2020). Learning improves decoding of odor identity with phase-referenced oscillations in the olfactory bulb. Elife 9, e52583.

Malinow, R., Schulman, H., and Tsien Richard, W. (1989). Inhibition of Postsynaptic PKC or CaMKII Blocks Induction But Not Expression of LTP. Science 245, 862–866.

Malvaut, S., Gribaudo, S., Hardy, D., David, L.S., Daroles, L., Labrecque, S., Lebel- Cormier, M.-A., Chaker, Z., Coté, D., De Koninck, P., et al. (2017). CaMKIIalpha Expression Defines Two Functionally Distinct Populations of Granule Cells Involved in Different Types of Odor Behavior. Current Biology 27, 3315–3329.e3316.

Martin, C., Beshel, J., and Kay, L.M. (2007). An olfacto-hippocampal network is dynamically involved in odor-discrimination learning. JNeurophysiol 98, 2196–2205.

Matsuo, N., Yamasaki, N., Ohira, K., Takao, K., Toyama, K., Eguchi, M., Yamaguchi, S., and Miyakawa, T. (2009). Neural activity changes underlying the working memory deficit in alpha-CaMKII heterozygous knockout mice. Front Behav Neurosci 3, 20.

Moberly, A.H., Schreck, M., Bhattarai, J.P., Zweifel, L.S., Luo, W., and Ma, M. (2018). Olfactory inputs modulate respiration-related rhythmic activity in the prefrontal cortex and freezing behavior. Nature Communications 9, 1528.

Mofleh, R., and Kocsis, B. (2021). Delta-range coupling between prefrontal cortex and hippocampus supported by respiratory rhythmic input from the olfactory bulb in freely behaving rats. Scientific Reports 11, 8100.

Nakazawa, K., McHugh, T.J., Wilson, M.A., and Tonegawa, S. (2004). NMDA receptors, place cells and hippocampal spatial memory. Nature Reviews Neuroscience 5, 361–372.

Namburi, P. (2021). Phase locking value. https://praneethnamburi.com/2011/08/10/plv/.

Nguyen Chi, V., Muller, C., Wolfenstetter, T., Yanovsky, Y., Draguhn, A., Tort, A.B., and Brankack, J. (2016). Hippocampal Respiration-Driven Rhythm Distinct from Theta Oscillations in Awake Mice. J Neurosci 36, 162–177.

Nolte, G., Bai, O., Wheaton, L., Mari, Z., Vorbach, S., and Hallett, M. (2004). Identifying true brain interaction from EEG data using the imaginary part of coherency. Clinical Neurophysiology 115, 2292–2307.

Purcell, S.M., Moran, J.L., Fromer, M., Ruderfer, D., Solovieff, N., Roussos, P., O’Dushlaine, C., Chambert, K., Bergen, S.E., Kahler, A., et al. (2014). A polygenic burden of rare disruptive mutations in schizophrenia. Nature 506, 185–190.

Robison, A.J. (2014). Emerging role of CaMKII in neuropsychiatric disease. Trends in Neurosciences 37, 653–662.

Rojas-Libano, D., Frederick, D.E., Egana, J.I., and Kay, L.M. (2014). The olfactory bulb theta rhythm follows all frequencies of diaphragmatic respiration in the freely behaving rat. Front Behav Neurosci 8, 214.

Scheffer-Teixeira, R., and Tort, A.B. (2016). On cross-frequency phase-phase coupling between theta and gamma oscillations in the hippocampus. eLife 5, e20515.

Siegle, J.H., and Wilson, M.A. (2014). Enhancement of encoding and retrieval functions through theta phase-specific manipulation of hippocampus. Elife 3, e03061.

Silva, A.J., Paylor, R., Wehner, J.M., and Tonegawa, S. (1992a). Impaired spatial learning in alpha-calcium-calmodulin kinase II mutant mice. Science 257, 206–211.

Silva, A.J., Rosahl, T.W., Chapman, P.F., Marowitz, Z., Friedman, E., Frankland, P.W., Cestari, V., Cioffi, D., Südhof, T.C., and Bourtchuladze, R. (1996). Impaired learning in mice with abnormal short-lived plasticity. Current Biology 6, 1509–1518.

Silva, A.J., Stevens, C.F., Tonegawa, S., and Wang, Y. (1992b). Deficient hippocampal long-term potentiation in alpha-calcium-calmodulin kinase II mutant mice. Science 257, 201–206.

Slotnick, B.M., and Restrepo, D. (2005). Olfactometry with mice. In Current Protocols in Neuroscience, J.N. Crawley, C.R. Gerefen, M.A. Rogawski, D.R. Sibley, P. Skolnick, and S. Wray, eds. (New York: John Wiley and Sons, Inc), pp. 1–24

Suarez-Pereira, I., Canals, S., and Carrion, A.M. (2015). Adult newborn neurons are involved in learning acquisition and long-term memory formation: the distinct demands on temporal neurogenesis of different cognitive tasks. Hippocampus 25, 51–61.

Tort, A.B., Komorowski, R., Eichenbaum, H., and Kopell, N. (2010). Measuring phase- amplitude coupling between neuronal oscillations of different frequencies. J Neurophysiol 104, 1195–1210.

Udakis, M., Pedrosa, V., Chamberlain, S.E.L., Clopath, C., and Mellor, J.R. (2020). Interneuron-specific plasticity at parvalbumin and somatostatin inhibitory synapses onto CA1 pyramidal neurons shapes hippocampal output. Nature Communications 11, 4395.

Vizcay, M.A., Duarte-Mermoud, M.A., and Aylwin Mde, L. (2015). Odorant recognition using biological responses recorded in olfactory bulb of rats. Comput Biol Med 56, 192–199.

Wang, P.Y., Boboila, C., Chin, M., Higashi-Howard, A., Shamash, P., Wu, Z., Stein, N.P., Abbott, L.F., and Axel, R. (2020). Transient and Persistent Representations of Odor Value in Prefrontal Cortex. Neuron 108, 209–224.e206.

Won, G.H., Kim, J.W., Choi, T.Y., Lee, Y.S., Min, K.J., and Seol, K.H. (2018). Theta- phase gamma-amplitude coupling as a neurophysiological marker in neuroleptic-naïve schizophrenia. Psychiatry Research 260, 406–411.

Yamasaki, N., Maekawa, M., Kobayashi, K., Kajii, Y., Maeda, J., Soma, M., Takao, K., Tanda, K., Ohira, K., Toyama, K., et al. (2008). Alpha-CaMKII deficiency causes immature dentate gyrus, a novel candidate endophenotype of psychiatric disorders. Molecular Brain 1, 6.

